# A global database of photosynthesis model parameters, and modelled photosynthetic responses from every major terrestrial plant clade

**DOI:** 10.1101/2020.10.06.328682

**Authors:** Mina Rostamza, Gordon G. McNickle

## Abstract

Plant photosynthesis is a major part of the global carbon cycle and climate system. Carbon capture by C_3_ plants is most often modelled using the Farquhar-von-Caemmerer-Berry (FvCB) equations. We undertook a global synthesis of all parameters required to solve the FvCB model. The publicly available dataset we assembled includes 3663 observations from 336 different C_3_ plant species among 96 taxonomic families coming from every major vascular plant clade (lycophytes, ferns, gymnosperms, magnoliids, eudicots and monocots). Geographically, the species in the database have distributions that span the majority of the globe. We used the model to predict photosynthetic rates for a hypothetical average plant in each major terrestrial plant clade and find that generally plants have dramatically increased their photosynthetic abilities through evolutionary time, with the average monocot (the youngest clade) achieving maximum rates of photosynthesis almost double that of the average lycophyte (the oldest clade). We also solved the model for different hypothetical average plant functional types (PFTs) and find that herbaceous species generally have much higher rates of photosynthesis compared to woody plants. Indeed, the maximum photosynthetic rate of graminoids is almost three times the rate of the average tree. The resulting functional responses to increasing CO_2_ in average hypothetical PFTs would suggest that most groups are already at or near their maximum rate of photosynthesis. However, phylogenetic analysis showed that there was no evidence of niche conservatism with most variance occurring within, rather than among clades (K=0.357, p=0.001). This high within-group variability suggests that average PFTs may obscure important plant responses to increasing CO_2_. Indeed, when we solved the model for each of the 3663 individual observations, we found that, contrary to the predictions of hypothetical average PFTs, that most plants are predicted to be able to increase their photosynthetic rates. These results suggest that global models should seek to incorporate high within-group variability to accurately predict plant photosynthesis in response to a changing climate.

## INTRODUCTION

Plant photosynthesis is a major factor in the global climate system. Indeed, the annual flux of atmospheric carbon (C) through the leaves of terrestrial plants is estimated to be 1 × 10^15^ g yr^−1^ (Beer *et al*., 2010, Hetherington & Woodward, 2003). Carbon capture by C_3_ plants is most often modelled using models derived from the Farquhar-von-Caemmerer-Berry (FvCB) equations (Farquhar *et al*., 1980, Farquhar & Wong, 1984, Sharkey *et al*., 2007, Von Caemmerer, 2000). The FvCB model is a process based physiological model that accurately describes the rate of photosynthesis across light levels, and across both CO_2_ and O_2_ concentrations. In its modern form, the FvCB model also accounts for triose phosphate limitation (Lombardozzi *et al*., 2018, Mcclain & Sharkey, 2019). Indeed, a version of the FvCB model forms the basis for most physiological, ecological, and earth system models that include plants (Rogers *et al*., 2017).

Models that incorporate plant photosynthesis require accurate parameter estimates, estimates which are spread across four decades of scientific inquiry and may be difficult to find for specific taxa. There have been several syntheses and meta-analyses that focus on two parameters of the FvCB model, *V*_*cmax*_ and *J*_*max*_ (E.g. Kattge & Knorr, 2007, Walker *et al*., 2014, Wullschleger, 1993), as well as syntheses on empirically estimated maximum photosynthetic rates (Gago *et al*., 2019), but we are unaware of any attempt at a global synthesis of the full suite of at least 12 parameters needed to fully predict photosynthetic rates across the all C_3_ plants. In addition, the modern FvCB model of photosynthesis is well known to be over-parameterised (Qian *et al*., 2012), and modern techniques for curve fitting and parameter estimation can benefit from better prior information. For example, Bayesian methods can work from a known prior distribution of parameter values to enhance the ability to accurately estimate parameters (e.g. Patrick *et al*., 2009). Thus, collecting all available parameter estimates into one database would greatly enhance the ability to model global photosynthesis, as well as our ability to estimate parameters for new taxa.

Here, we describe a synthesis of all FvCB parameters where at least one parameter was estimated for a given species. The summary includes parameter estimates from 359 different plant species from 96 taxonomic families coming from every major vascular plant clade (lycophytes, ferns, gymnosperms, magnoliids, eudicots and monocots) whose distributions span the majority of the globe. The parameter estimates are presented using a number of summary statistics and probability density histograms. We also solve the FvCB model using the full range of parameter estimates to generate predictions about the breadth of plant photosynthetic responses across major vascular plant clades, plant functional types, and individual leaves. The full dataset containing 3663 unique rows of data is publicly available.

## MATERIALS AND METHODS

### The FvCB photosynthesis model

Here, we briefly describe the equations of the FvCB model we used to seek parameterizations (Farquhar *et al*., 1980, Sharkey *et al*., 2007, Von Caemmerer, 2000). The most basic modern FvCB photosynthesis approach for C_3_ plants assumes that the rate of carbon assimilation (*A*) in photosynthesis is co-limited by either carbon (*A*_*c*_), light (*A*_*j*_) or TPU (*A*_*p*_) according to:

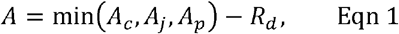

where *R*_*d*_ is the daytime respiration rate (See Table 1 for units).

**TABLE 1:** Summary of symbols, associated equation numbers in the text, plain descriptions and units for the parameters of the FvCB photosynthesis model that are included in this global summary. Note that those parameters in units of pressure (Pa), are reported differently by many authors. See text for unit conversions. Note that μmol m^-2^ s^-1^ represents carbon or photons depending on parameter.

| Parameter | Units | Eqn | Description |
| --- | --- | --- | --- |
| $V_{c,max}$ | $\mu\text{mol m}^{-2} \text{s}^{-1}$ | 2 | The carbon saturated maximum rate of carbon fixation. |
| $J_{max}$ | $\mu\text{mol m}^{-2} \text{s}^{-1}$ | 4, 5, 6 | The light saturated maximum possible rate of electron transport. |
| $C_i$ | Pa | 2, 3 | Intercellular partial pressure of $\text{CO}_2$ at the site of RUBISCO activity. (When constructing a $\text{CO}_2$ response curve, this is considered a variable, not a parameter.) |
| $\Gamma^*$ | Pa | 2, 3 | The partial pressure of the $\text{CO}_2$ compensation point. This is equivalent to a giving up density. It is where carbon assimilation (i.e. benefits) equal respiration costs. |
| $K_c$ | Pa | 2 | The half saturation constant for $\text{CO}_2$ of RUBISCO. This is sometimes called the enzyme "affinity" for the substrate, though in reality it has limited biological meaning. Rather, it describes the the curvature of the $\text{CO}_2$ functional response. It is simply the partial pressure that is half-way to the maximum rate. |
| $K_o$ | Pa | 2 | The half saturation constant for $\text{O}_2$ of RUBISCO. This is sometimes called the enzyme "affinity" for the substrate, though in reality it has limited biological meaning. Rather it describes the curvature of the oxygenation functional response of RUBISCO. It is simply the partial pressure of oxygen that is half-way to the maximum rate. |
| $R_d$ | $\mu\text{mol m}^{-2} \text{s}^{-1}$ | 1 | The leaf respiration rate during the day of the leaf. |
| $T_p$ | $\mu\text{mol m}^{-2} \text{s}^{-1}$ | 7a, b | The rate of TPU production. |
| $r$ | unitless | 7a | The rate of TPU recycling. Often assumed to be 0. |
| $\alpha$ | unitless | 4 | The efficiency of light conversion. |
| $\theta$ | unitless | 4 | A curvature factor of the light response. |
| $a$ | unitless | 5 | The efficiency of light conversion. |
| **Variable** |  |  |  |
| $Q$ | $\mu\text{mol photons m}^{-2} \text{s}^{-1}$ | 4, 5 | The light photon flux density striking the leaf. (When constructing a $\text{CO}_2$ response curve, this can be considered a parameter not a variable) |
| $J$ | $\mu\text{mol m}^{-2} \text{s}^{-1}$ | 3, 4, 5 | The actual rate of electron transport going to support $\text{NADP}^+$ reduction for RuBP regeneration. |
| $A_c$ | $\mu\text{mol m}^{-2} \text{s}^{-1}$ | 1 | Rate of carbon assimilation limited by carbon reactions. |
| $A_j$ | $\mu\text{mol m}^{-2} \text{s}^{-1}$ | 1 | Rate of carbon assimilation limited by light and electron transport. |
| $A_p$ | $\mu\text{mol m}^{-2} \text{s}^{-1}$ | 1 | Rate of carbon assimilation limited by TPU. |
| $A$ | $\mu\text{mol m}^{-2} \text{s}^{-1}$ | 1 | Actual rate of carbon assimilation |

The carbon-limited portion of assimilation by photosynthesis in Eqn 1 is given by:

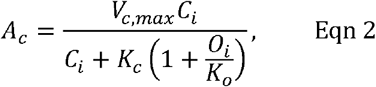

Where *C*_*i*_ and *0_i_* are the intracellular concentrations of CO_2_ and O_2_ at the site of ribulose-1,5-bisphosphate carboxylase/oxygenase (RuBisCO) activity respectively; *K*_*c*_ and *K*_0_ are the RuBisCO half saturation constants for CO_2_ and O_2_, respectively, and; *V*_*cmax*_ is the maximum possible rate of photosynthesis. Half saturation constants are often called the enzyme “affinity” for the substrate, but in reality they have limited biological meaning and simply describe the shape of the curvature of the functional response (Mcnickle & Brown, 2014).

The light limited portion of photosynthesis is given by:

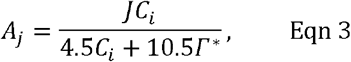

where *J* is the realised electron chain transport determined by light; *Γ\** is the minimum partial pressure of CO_2_ where carbon assimilation balances respiration (i.e *A* = *R*_*d*_) generally called the CO_2_ compensation point; and *C*_*i*_ is as above.

For the purposes of a global summary, the variable has *J* been determined in several ways over the years. The most common approach following Farquhar and Wong (1984) *J* was found by solving for the root of a simple quadratic equation:

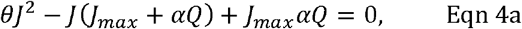

Where *Q* is the light photon flux density striking the leaf (μmol photons m^-2^ s^-1^), *J*_*cmax*_ is the light saturated maximum possible rate of electron transport; *θ* represents curvature of the light response; and; *α* is the efficiency of light conversion. Since this is a quadratic equation, and since negative values of *J* have no biological meaning, we can use the quadratic formula to find the positive root of Eqn 4a where:

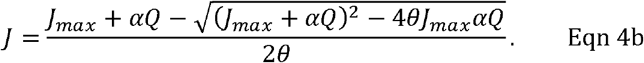

For some reason, most modern papers use different symbols from the original formulation. In addition, though significantly less common (Buckley & Diaz-Espejo, 2015), some authors use an approximation of eqns 4 where *J* is approximated according to:

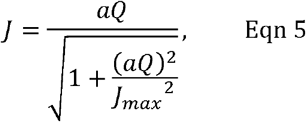

Where *a* represents the efficiency of light conversion (similar to α in eqn 4), and *Q* and *J*_*max*_ have their usual meanings. Note that *a* in Eqn 5 is more typically written in the literature as the Greek letter ‘*alpha*’, but we altered this to avoid confusion with Eqn 4.

Finally, TPU limitation of photosynthesis in its most detailed form is given by:

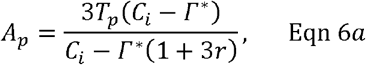

where *r* is a unitless scalar related to the proportion of glycolate recycled in chloroplasts (where 0 < *r <* 1);*T*_*p*_ is the rate of TPU, and the other parameters have their usual meanings given above (Table 1). However, it appears to be most common to assume that no glycolate is recycled (i.e. *r* = 0), and then Eqn 6a simplifies to:

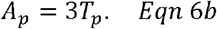

We note that there are even more complex versions than we have detailed here in Eqns 1-6. These more complex versions may include parameters for stomatal conductance of CO_2_ (Collatz *et al*., 1992), mesophyll conductance of CO_2_ (Flexas *et al*., 2014, Niinemets *et al*., 2011) and transpiration (Farquhar & Sharkey, 1982, Farquhar & Wong, 1984). However, leaf resistance to CO_2_ and transpiration could be considered their own rather large sub-fields independent from many attempts to estimate FvCB model parameters. Thus, we purposefully omitted resistances from our survey of the literature to make the scope of the literature synthesis more tractable. Also, models that include detailed temperature relationships via the Arrhenius equation are also somewhat common (Kattge & Knorr, 2007). Again however, to limit the scope of our synthesis we did not collect the parameters of these Arrhenius rate parameters for a more complex temperature dependent FvCB model. We view these as future updates that could be made to the dataset.

### Literature search

Briefly, we began with a search for “Farqhuar photosynthesis models” in the Web of Science, read each paper carefully, and extracted values for all parameters of the FvCB model (Table 1). The search was expanded by using the bibliographies of these papers. In total data was extracted from 202 papers. Units were standardized across studies for comparison. We also collected as many methodological details about the conditions in the cuvette as possible to attempt to understand sources of variation in the data that were not caused by taxonomy (E.g. humidity, temperature, vapour pressure deficit, light levels, CO_2_ pressure). The full details of our systematic literature search and data extraction methods and criteria are given in the Supplementary Information.

We found that studies could generally be broadly organised into three types: temperature effects, leaf nitrogen effects, or mean plant effects. First, temperature is well known to affect the biochemical reactions in photosynthesis and we wanted to account for this variability (Kattge & Knorr, 2007). Thus, when temperature was the treatment parameter, values at each temperature were recorded separately. These rows are labelled as type ‘*T*’ in the data file. Second, leaf nitrogen can also be related to photosynthesis parameters and we wanted to account for this variability as well (Kattge *et al*., 2009, Walker *et al*., 2014). Thus, when parameters were estimated separately by leaf nitrogen, each leaf nitrogen level was recorded separately. Nitrogen was sometimes reported as percent dry weight, and sometimes as mass per leaf area. Either were recorded, but in separate data columns. These rows are labelled as type ‘*N*’ in the data file. Studies that reported just one value for each parameter and species (i.e. a species mean) are labelled as type ‘*M*’ in the data file. Thus, there are three types of data: (i) mean values (type *M*); (ii) leaf temperature manipulations (type *T*), and; (iii) leaf nitrogen differences (type *N*). However, when available, temperature was recorded for type *M* and type studies. Similarly, when available, leaf nitrogen was recorded for type *T* and type *M* studies.

### Taxonomy and biogeography

Species names reported in the original papers were checked the National Center for Biotechnology Information’s species taxonomy database using the brranching::phylomatic_names function in R (Webb & Donoghue, 2005). In general, this just updated any outdated species names to the most modern accepted name. In one case, we had to manually change a species names to a taxonomic synonym to match to the database: *Echinochloa crus*-galli (L.) Beauv was changed to an accepted synonym *Digitaria hispidula* (Retz.) Willd. The updated species names were then pruned from the plant megatree of Zanne *et al*. (2014) to visually represent the taxonomic coverage of the dataset using the brranching library in R (v. 0.6; Chamberlain, 2020). The literature search produced a few parameter estimates for 21 C4 graminoid species that others had reported in the literature and the FvCB model is only appropriate for C_3_ photosynthesis (Collatz *et al*., 1992). The final data file includes these few C4 parameters, but C4 plants are excluded from all analyses described here because the FvCB model does not apply to them. Species were also assigned to the following broad plant functional types (PFTs) based on growth form: C_3_ graminoid, Forb, Vine/Climbing, Shrub, or Tree (Also, C4 graminoids are labelled in the datafile, but we do not analyse them here). Finally, we recorded growth habit of each species in the following categories: annual, biennial, perennial, or crop for herbaceous species; deciduous, evergreen, or crop for woody species; and fern, tree, or club moss for ferns. Growth habit information was not available for three rare and exotic species and was recorded as NA. Of course, future users are free to organize species into whatever other categories are of interest.

Not all studies reported a location of plant material collection, measurement, or the cultivar examined. Thus, to obtain a sense of the geographic coverage of species in this dataset, we used the Botanical Information and Ecology Network (BIEN; Maitner *et al*., 2018) to obtain museum occurrence records for each species in our global database. We then mapped these occurrence records. The resulting map shows the global distribution of all species in our dataset. Importantly this map is not the global distribution of measurement locations. Rather, the resulting map provides window into the range distributions of all species that have been studied for photosynthesis, and therefore details what regions of the world have good coverage of at least approximate species level photosynthetic data. In addition, the resulting map also shows what regions would benefit from increased empirical attention to improve global models.

### Data summary

To create a global summary table, we treated the three data types differently. For studies that report only mean values (type “M”), we used all values in the summary. For temperature studies (type “T”), we only used the value nearest to 25°C. This was done because 25°C was the most common temperature used in studies that did not manipulate leaf temperature and most temperature studies measure the same leaf across many temperatures. Our approach of using only one value per leaf was to avoid pseudoreplication. For leaf nitrogen studies (Type “N”), we averaged values across leaf N amounts to capture the mean response of plants growing across different soil fertilities. Because different leaf nitrogen contents represented individual plants in each study, this method creates a species mean and also seeks to avoid pseudoreplication. To generate a global summary, we calculated the mean, standard deviation, median, maximum, minimum, skew, and kurtosis for each parameter from Eqns 1-6.

Data were also summarised by major taxonomic clade and PFT. For these summary rows, there were generally too few categories in most groups to create density distributions, and we report only the mean, standard deviation, and sample size. For sample sizes that were n<3, we report the standard deviation as NA. For these summaries, we did not separate the *T, M*, and *N* data types because many taxa were only studied once.

### Breadth of model outcomes

It is useful to the modelling community to have a large database of FvCB model parameters, but the raw parameters do not show the breadth of plant photosynthetic responses on their own, since 12 parameters (Table 1) may combine in a many different ways to produce the same rate of photosynthesis. To examine the range of predicted photosynthetic responses, we used the parameter estimates to actually solve the FvCB model for plant carbon assimilation rate by solving for model predicted *A − C_i_* and *A−Q* curves. We did this in three ways. All three of these analysis uses all three types of data (M, T and N) in order to show the breadth of responses.

First, we solved the model for the average hypothetical plant that describes each major plant clade. To do this, each parameter value was set equal to the average of the clade, and the predicted*A − C_i_* and *A−Q* curves were solved. In addition, all parameters were set to the upper and lower 95% confidence intervals around the mean, and the equations were solved again to give a sense of variation within the clade. Missing values were replaced with the global mean in the clade analysis. This let us compare the average photosynthetic functional response of some hypothetical average lycophyte, fern, gymnosperm, magnoliid, eudicot, and monocot.

Second, we solved the model for the average hypothetical plant that describes each PFT. This was done as above with the mean parameter values and the upper and lower confidence intervals. Here, missing values were replaced with the mean of the appropriate clade (e.g. missing graminoid parameters were replaced with the monocot mean, while missing vine or shrub parameters with the eudicot mean).

Third, we went down each of the 3663 rows of our dataset and solved the model for every individual leaf for which we had data. However, no study in our synthesis estimated all parameters of the FvCB equation. Thus, to fill in gaps for any row, we used the global mean value for any missing parameter values. In addition, the average *A*_*max*_ for each species was calculated from these model runs and then drawn onto the phylogeny. We used Bloomberg’s K to examine phylogenetic signal in the species level data using phytools in R (Revell, 2012).

## RESULTS

### Taxonomy and biogeography

In total, we obtained at least one parameter estimate from 359 species in 96 families (Fig 1A, Table S1). The data included all major vascular plant clades including: lycophytes (2 species, 1 family), ferns (33 species, 16 families), gymnosperms (23 species, 3 families), and angiosperms (303 species, 77 families). Angiosperms can be further separated into three more sub-clades comprised of monocots (62 species, 7 families), eudicots (235 species, 68 families), and magnoliids (6 species, 3 families). Tables S1 and S2 include more detailed summaries breaking the available data up among each of the 96 taxonomic families and within the six clades. In addition, Tables S3 and S4 contain more detailed summaries breaking the data up by PFT and growth habit.

**FIG 1:**
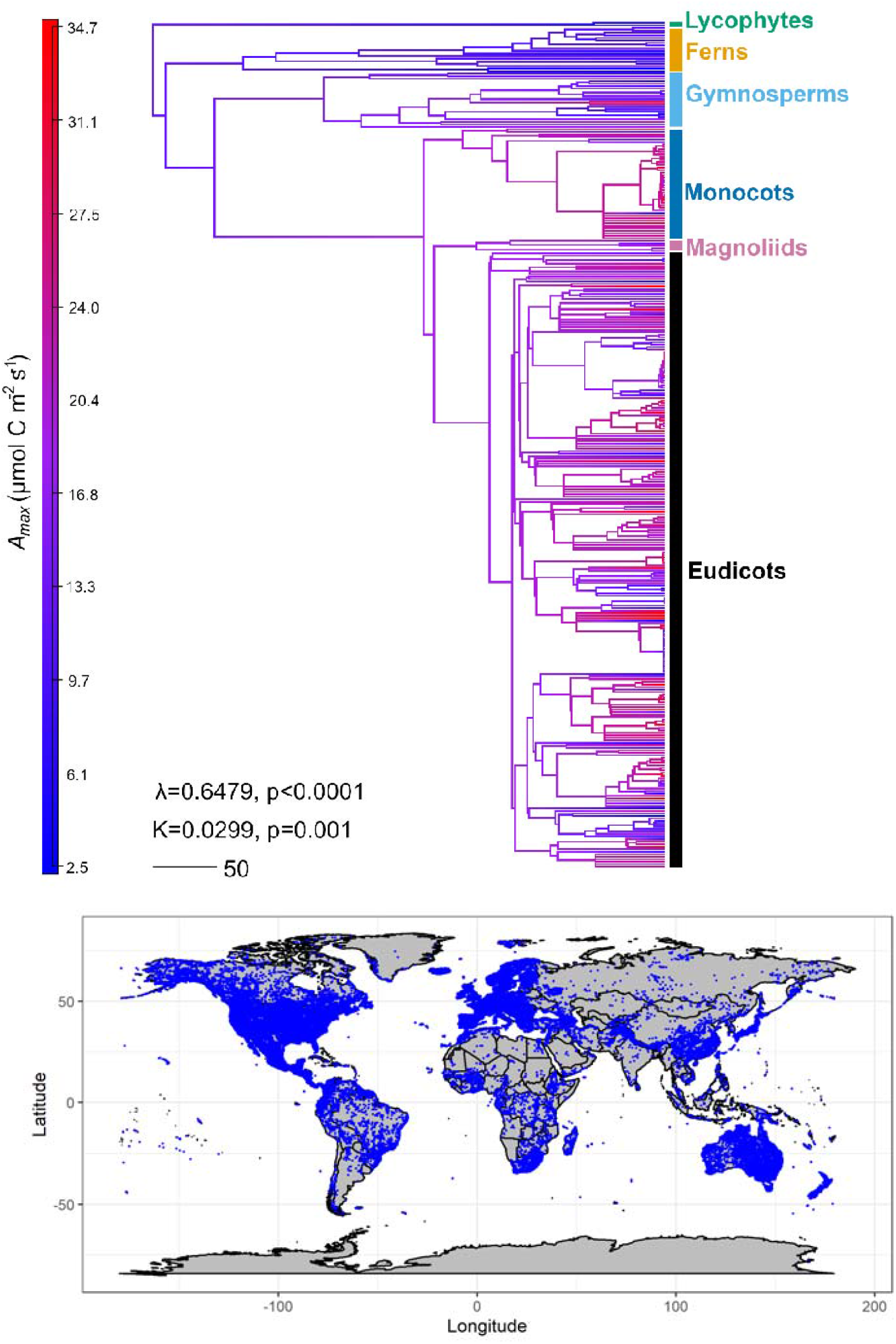
(A) The evolutionary relationships among all 359 species among 6 major terrestrial plant clades for which photosynthesis parameters were recovered from the literature and included in our database. Scale bar represents 50 million years of evolution, and colours represent modelled. Unresolved taxonomy are drawn as polytomies at the genus level. The raw phylogeny in newick format is given in the supplementary information to allow digital visualisation of such a large phylogeny. There was no evidence of phylogenetic niche conservatism across the entire dataset, meaning species with low or high are equally likely in any clade. (B) Global distribution data for all 334 species included in the data set (blue points). Points represent occurrence distribution data for all species retrieved from BIEN, not the locations of the measurements.

The occurrence data from BIEN shows that the majority of data comes from North America, Europe, Australia, New Zealand, and Japan (Fig 1B). Even though much of Africa is desert containing few to no plants, coverage on the vegetated parts of the continent was patchy, with the best coverage in South Africa and parts of west Africa. However, excluding regions in Africa that mostly do not contain plants (i.e. the Sahara and Namib Deserts), there are entire countries in southern Africa for which very few species have been studied (e.g. Angola, Morocco, Nigeria). Similarly, coverage was spotty in northern Asia (E.g. Russia, Kazakhstan, Mongolia), Indochina, Indonesia and South America (e.g. Chile, Argentina, Uruguay, Paraguay). This is problematic, because many of these regions of Africa, Asia, and South America with patchy data are known diversity hotspots where a small number of taxonomic samples may not represent the average plant in those regions (Myers *et al*., 2000). There was, however, a surprising amount of data from species endemic to the southern foothills of the Himalayan mountains, and from Colombia and Ecuador. Regions of Brazil’s Amazon rainforest are also sparsely measured, with much of the Brazilian data appearing to have come from the southern grassland regions.

Ignoring national borders, the geographic coverage suggests that most named ecosystem ‘types’ (e.g. sensu Whittaker, 1975) also have good coverage. Temperate and boreal forests and grasslands have particularly good coverage (Fig 1B). However, given the high species diversity of tropical ecosystems, there are likely important gaps in our understanding of the diversity of photosynthetic responses in tropical regions across the globe. The arctic regions of Europe have very good coverage, but there is little data from arctic Russia and large gaps in coverage for arctic North America. Similarly, for grasslands, there is very good coverage in Australia and North America, but relatively little for the grasslands of Asia, Africa, and South America.

### Summary statistics and parameter distributions

By far, *V*_*cmax*_ (n=1364), *J*_*max*_ (n=961), and, to a lesser extent,*T*_p_ (n=171) were the most frequently estimated and reported parameter values (Table 2, 3). In general, most of the probability density distributions of observed parameters were skewed or possibly multi-modal (Table 2, 3, Fig S1, Fig S2). It is noteworthy that the minimum and maximum reported values of most parameters differed by as much as four orders of magnitude across all vascular plants. Also, the coefficient of variation in most cases was 0.5 or higher, suggesting high dispersion of the data among species. However, the data show that the majority of this within- and among-species variation over orders of magnitude was driven by methodological differences among research groups (Supplementary information). For example, a few species were studied many times by different groups, and variation in reported parameters for these species were largely explained by the nitrogen content of the leaves each group measured, and the VPD tolerance they used when taking measurements (Table S5, Fig S3). Similarly, among all species in our data base, methodological choices of different research groups explained 60% of the variation in parameter estimates (Fig S4, Fig S5). Thus, once methods are controlled, the variation is primarily driven by species differences.

**TABLE 2:** Summary statistics and distribution parameters for each parameter of the Farquhar photosynthesis equations (Eqn 1-4). The global density distribution histogram for each parameter is shown in Fig S1. The mean, standard deviation (SD) and sample size is also shown for each major clade. SD is reported as NA for n<3, and empty cells represent no data.

| Clade | Stat | $V_{c \max}$<br>($\mu\text{mol m}^{-2} \text{s}^{-1}$) | $J_{\max}$<br>($\mu\text{mol m}^{-2} \text{s}^{-1}$) | $C_i$<br>(Pa) | $\Gamma^*$<br>(Pa) | $K_c$<br>(Pa) | $K_o$<br>(Pa) | $R_d$<br>($\mu\text{mol m}^{-2} \text{s}^{-1}$) | $T_p$<br>($\mu\text{mol m}^{-2} \text{s}^{-1}$) |
| --- | --- | --- | --- | --- | --- | --- | --- | --- | --- |
| All | n | 1352 | 953 | 88 | 58 | 43 | 29 | 458 | 171 |
|  | Mean | 61.0 | 134.8 | 37.8 | 4.5 | 35.8 | 29169.1 | 1.2 | 12.0 |
|  | SD | 39.1 | 81.0 | 6.6 | 2.6 | 12.6 | 9829.1 | 1.7 | 5.4 |
|  | CV | 0.6 | 0.6 | 0.17 | 0.58 | 0.35 | 0.33 | 1.4 | 0.45 |
|  | Median | 52.9 | 118.0 | 31.8 | 4.0 | 30.5 | 28168.4 | 0.8 | 12 |
|  | Min | 0.3 | 10.4 | 15.9 | 0.01 | 17.0 | 16500 | 0.02 | 1.7 |
|  | Max | 267.1 | 498.4 | 37.7 | 9.2 | 70.7 | 47926.7 | 20.3 | 29.2 |
|  | Skew. | 1.09 | 1.09 | -0.64 | 0.18 | 0.82 | 0.43 | 7.62 | 0.64 |
|  | Kurtosis | 4.60 | 4.41 | 2.01 | 2.45 | 3.08 | 2.04 | 75.31 | 3.59 |
| Lycoph. | n | 2 | 2 |  |  |  |  | 2 |  |
|  | Mean | 17.2 | 25.5 |  |  |  |  | 0.5 |  |
|  | SD | NA | NA |  |  |  |  | NA |  |
| Ferns | n | 33 | 33 |  |  |  |  | 33 |  |
|  | Mean | 51.2 | 59.7 |  |  |  |  | 0.5 |  |
|  | SD | 27.9 | 24.1 |  |  |  |  | 0.3 |  |
| Gymno. | n | 461 | 301 | 37 | 27 | 2 | 2 | 41 | 49 |
|  | Mean | 34.7 | 65.2 | 19.4 | 7.7 | 27.4 | 41543.3 | 0.9 | 2.8 |
|  | SD | 32.25 | 42.93 | 5.14 | 3.04 | NA | NA | 0.45 | 2.62 |
| Magno. | n | 18 | 3 |  |  |  |  | 1 |  |
|  | Mean | 21 | 119.3 |  |  |  |  | 0.7 |  |
|  | SD | 12.85 | 29.16 |  |  |  |  | NA |  |
| Eudic. | n | 2037 | 1060 | 43 | 63 | 42 | 31 | 283 | 48 |
|  | Mean | 76.3 | 135.9 | 28.8 | 4.3 | 47.5 | 26442.6 | 1.5 | 6.8 |
|  | SD | 56.88 | 64.8 | 5.67 | 1.87 | 31.63 | 11382.05 | 1.76 | 2.6 |
| Monoc. | n | 395 | 342 | 15 | 8 | 22 | 20 | 225 | 236 |
|  | Mean | 86.6 | 196.7 | 28.8 | 3.0 | 51.9 | 37944 | 0.79 | 13 |
|  | SD | 40.9 | 92.8 | 7.0 | 1.7 | 32.59 | 17120.35 | 0.50 | 5.35 |
| **PFT** |  |  |  |  |  |  |  |  |  |
| C3<br>gramin. | n | 388 | 342 | 12 | 5 | 21 | 20 | 222 | 236 |
|  | Mean | 87.2 | 199.6 | 29.7 | 3.0 | 53.6 | 37944 | 0.8 | 13 |
|  | SD | 41.1 | 92.2 | 7.1 | 1.7 | 32.8 | 17120.3 | 0.5 | 5.4 |
| Forb | n | 222 | 123 | 22 | 43 | 23 | 21 | 40 | 6 |
|  | Mean | 83.5 | 167.5 | 31.2 | 4.3 | 41.1 | 24574.4 | 1.5 | 9.4 |
|  | SD | 50.3 | 75.4 | 6.0 | 1.6 | 32.2 | 10685.8 | 2.2 | 3.4 |
| Vine | n | 55 | 54 | 1 | 1 |  |  | 5 | 2 |
|  | Mean | 55.9 | 95.4 | 20 | 4.1 |  |  | 0.5 | 9.1 |
|  | SD | 23.1 | 37.2 | NA | NA |  |  | 0.3 | NA |
| Shrub | n | 166 | 143 | 3 | 4 | 3 |  | 23 | 21 |
|  | Mean | 72.1 | 125.7 | 18.7 | 2.8 | 26.8 |  | 2.1 | 6.6 |
|  | SD | 58.4 | 63.3 | 0.8 | 3.5 | 5.0 |  | 1.7 | 2.5 |
| Tree | n | 2077 | 1048 | 57 | 42 | 19 | 12 | 259 | 68 |
|  | Mean | 66.6 | 114.4 | 22.4 | 6.6 | 54.9 | 32073 | 1.3 | 3.7 |
|  | SD | 56.1 | 65.4 | 6.0 | 3.0 | 30.1 | 12035.2 | 1.6 | 2.7 |

\* only $R_d$ values reported with estimates of FvCB model parameters were sought. This will not be a complete literature review of plant respiration rates.

**TABLE 3:**
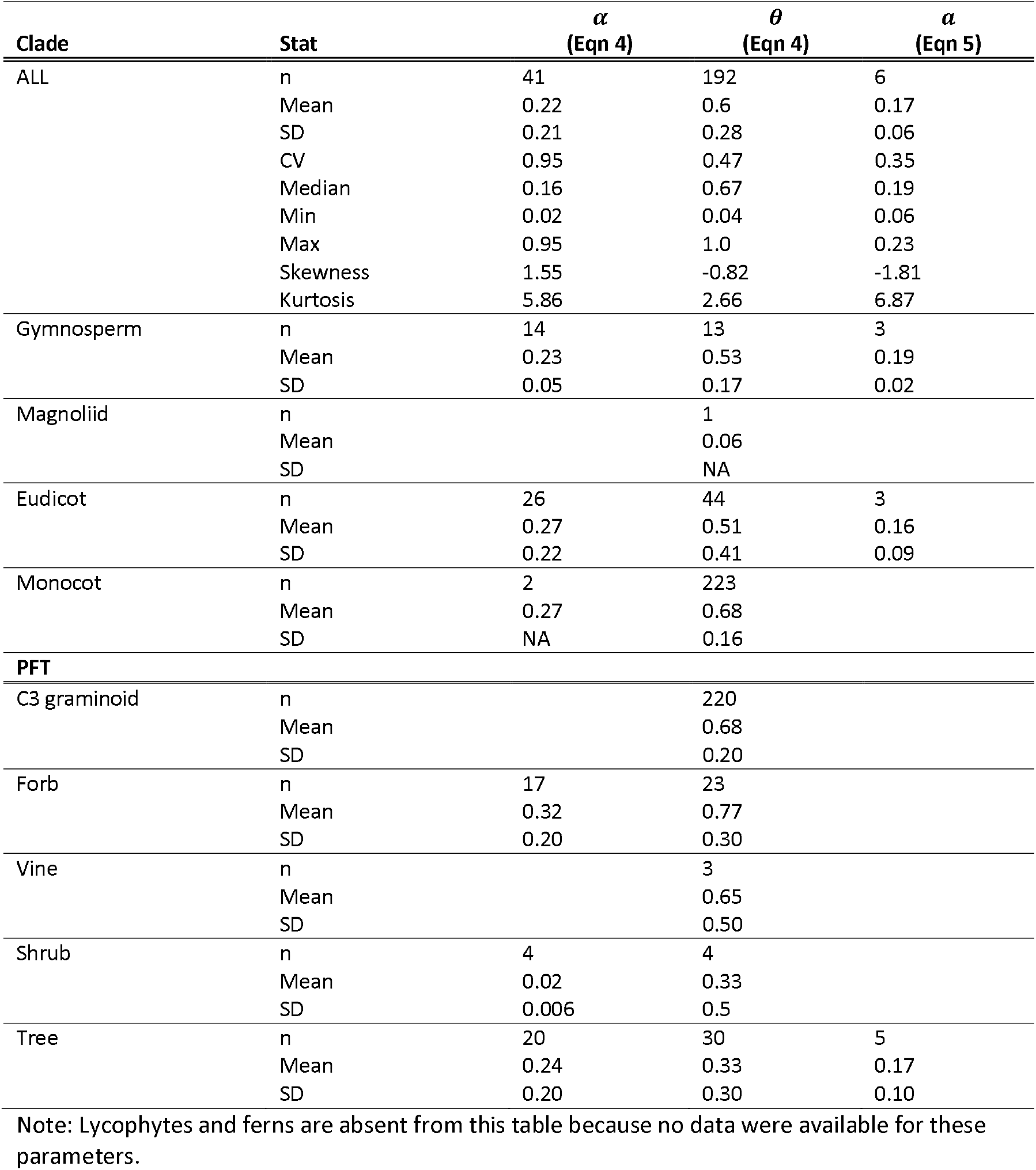
Summary statistics across all measurements for 334 species for each parameter of the Farquhar photosynthesis equations 4-5. The global density distribution histogram for each parameter is shown in Fig S2. The mean, standard deviation (SD) and sample size is also shown for each major clade. SD is reported as NA for n<3, and empty cells represent no data. Clades are summarised by family in Table S1 and by functional type growth habit in Table S4.

Among the six major clades, eudicots had the most data (n=2037 for *V*_*cmax*_) followed by gymnosperms (n=461 for *V*_*cmax*_) and monocots (n=400 for *V*_*cmax*_). In general, for both *V*_*cmax*_ and *J*_*max*_ there was a clear order to the means such that monocots > eudicots > magnoliids > gymnosperms > lycophytes. This suggests that plants have become increasingly adept at photosynthesis through evolutionary time. However, ferns do not seem to fit into this schema with ferns in-between magnoliids and gymnosperms. The half saturation constants (*K*_*c*_ and *K*_*0*_) were only estimated twice for gymnosperms and never for lycophytes, ferns, and magnoliids. Better in vivo sampling of lycophyte, fern and magnoliid species may resolve phylogenetic differences of parameter values in the future.

For PFTs, most researchers studied trees (n=2077 for *V*_*cmax*_) followed by C_3_ graminoids (n=388 for *V*_*cmax*_) and forbs (n=222 for *V*_*cmax*_). There were relatively few vines, and shrubs. When they are included there was a clear order, for both *V*_*cmax*_ and *J*_*max*_ means such that C_3_ graminoids > forbs > shrubs > trees > vines.

### Model outcomes

We also used the parameters to solve the FvCB model to explore predicted photosynthesis rates. For comparison among the six major clades represented in our data, we note that *A*_*max*_ was significantly different such that monocots > eudicots > gymnosperms (Fig 2A, B). There were too few parameter estimates to draw confidence intervals for lycophyte, fern, and magnoliid clades, but readers should assume they are very wide, and we hesitate to draw many conclusions without more data. However, if the patterns stand, these predicted photosynthetic rates among clades show a similar pattern to empirically estimated photosynthetic rates showing that on average land plants have substantially increased their photosynthetic ability over evolutionary time, matching empirical observations of *A*_*max*_ (Gago *et al*., 2019).

**Fig 2:**
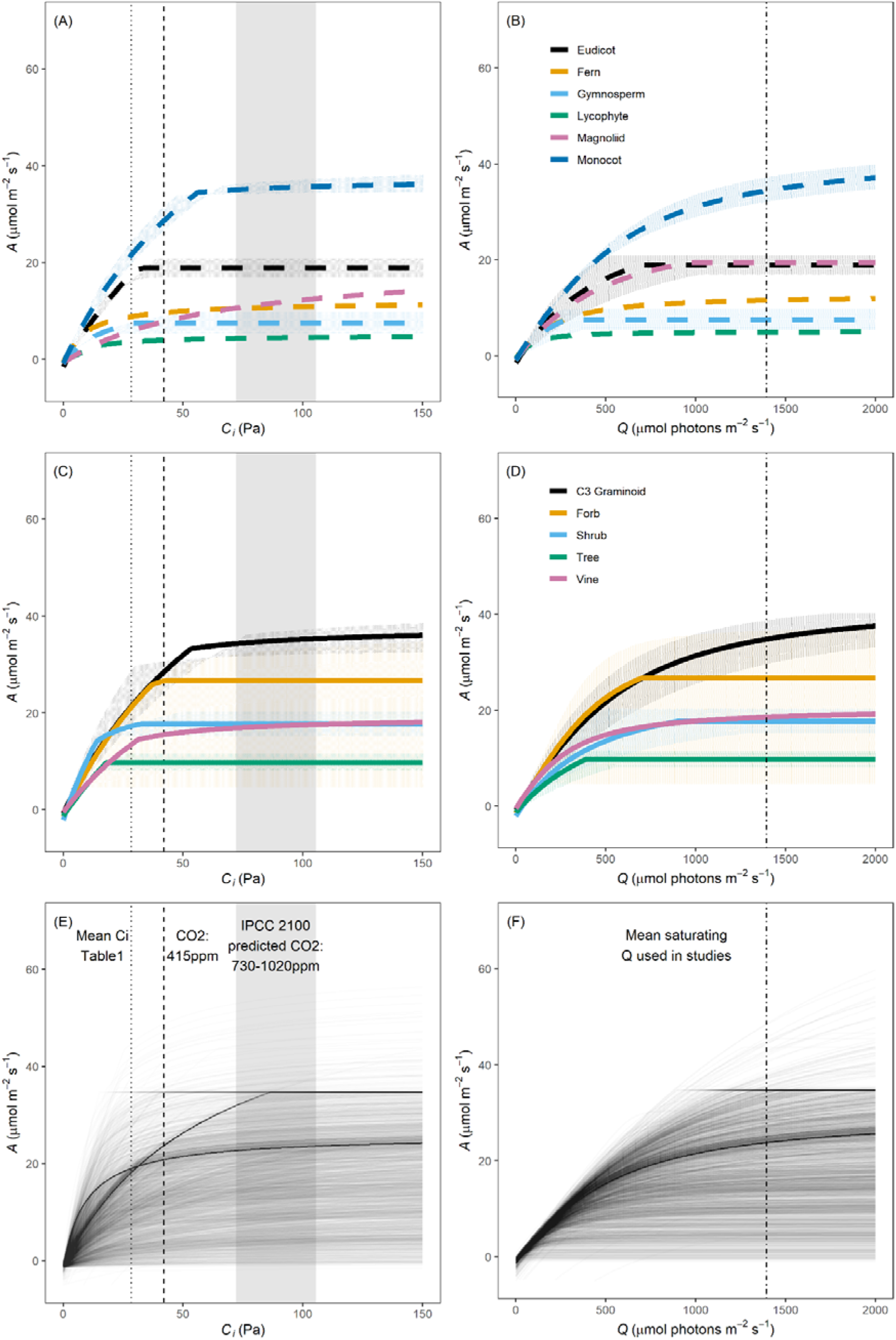
(**A**) Mean *A − C_i_* and (**B**) *A−Q* curves among each of the six major plant clades for which we have parameter estimates (solid lines). We also show (**C**) Mean *A − C_i_* and (**D**) *A−Q*curves with species grouped by broad functional type (dashed lines). Shading around the mean curve in panels A-D represents solving the model with the 95% confidence interval around all parameters. These 95% confidence intervals cross on the *A − C*_i_ curve because of the oxygenation behaviour of RUBISCO at low CO_2_ partial pressures. For, lycophytes, ferns, magnoliids, and vines the large number of missing values were replaced with the global mean, and we did not draw confidence intervals for these groups (readers should assume they are very wide due to low taxonomic sampling). Finally, each line in the lower panels represent 3663 separate (**E**) *A − C*_i_ or (**F**) *A−Q* curves generated for every single row in our database and show the breadth of individual species responses across diverse conditions. For these 3663 lines, missing values were replaced with the global mean (Table 2). In all panels, for *A − C*_i_ curves, the black vertical dotted line marks the estimated mean partial pressure of CO_2_ inside the leaf in reported in the literature, while the dashed line represents the approximate current partial pressure of CO_2_ in the atmosphere at the time of writing. The shaded rectangle represents the predicted range of CO_2_ Intergovernmental panel on climate change (IPCC) predicted partial pressures of CO_2_ by the year 2100. For *A−Q* curves, the black dot-dash line represents the mean saturating Q used among studies in our database.

For PFT photosynthetic rates, *A_max_* was significantly different such that C_3_ graminoids > shrubs > trees, but the confidence interval around forbs was so large that this group was not significantly different from any of these groups, suggesting herbaceous forbs fill a wider variety of photosynthetic niches than other groups (Fig 2C,D). Like magnoliids, there were too few parameter estimates for vines to draw confidence intervals for these PFTs, and we hesitate to draw conclusions about these PFTs.

There is reason to think analysing some hypothetical mean clade or PFT member might obscure diverse responses of genotypes and species. Thus, we also went down our dataset row by row to generate the entire breadth of photosynthetic responses of every leaf and species for which we had data. When parameters were missing, we replaced them with the global mean (Table 1). This created 3663 *A − C_i_* and 3663 *A−Q* curves (Fig 2E,F). When all 3663 curves were plotted on the same axis but with slightly transparent lines, the darker regions show the most common responses and the lighter regions show fewer common responses. Not surprisingly, the darkest lines largely appear to trace the average monocot and eudicot because these were best taxonomically sampled groups. The breadth of maximum photosynthesis values ranged from slightly negative rates of photosynthesis (−1.6 μmol C m^-2^ s^-1^) to 59.7 μmol C m^-2^ s^-1^.

When these data were averaged by species, there was no phylogenetic niche conservatism (Fig 1A, K=0.357, p=0.001) indicating that the majority of variation occurred within clades rather than among clades. Therefore, even though clades do appear to differ on average (Fig 2A,B), each clade is just as likely to contain some individual species with either low or high modelled *A*_*max*_ (Fig 1A).

## DISCUSSION

Photosynthesis is an important part of the earth climate system. Enormous quantities of CO_2_ pass through terrestrial plants each year across the globe. As such, accurate parameterisation of photosynthetic models is key to an understanding of phenomena ranging from simple leaf level physiology to global atmospheric dynamics. Here, we undertook a synthesis of parameter values in the literature. The data set we assembled from the literature includes 3663 rows of data across 359 different plant species from 96 taxonomic families that spanned all major vascular plant clades including lycophytes, ferns, gymnosperms, magnoliids, monocots, and eudicots (Fig 1; Table 2). Our aim was that modellers with diverse interests and questions could make use of this dataset, either for some general plant summarised by clade or PFT (Table 2,3). We also hope that empirical estimations of new parameter estimates can benefit from detailed prior probability distributions for modern Bayesian model fitting approaches (Fig S1, S2). However, it is also possible with these data to get more detailed, for example by family (Table S1, Table S2), growth habit (Table S3, Table S4), or even by individual species.

It has been shown that empirical estimates of *A*_*max*_ show a trend towards increasing photosynthetic efficiency in C_3_ plants through evolutionary time such that lycophytes > ferns > gymnosperms > angiosperms (Gago *et al*., 2019). It is exciting that the FvCB model is accurate enough to return the same ranking when estimated parameters are used to predict *A*_*max*_ (Fig 2). It’s not surprising that increases in are due to evolutionary adaptations that lead to increases in both *V*_*max*_ and *V*_*max*_ since these parameters control the maximum photosynthetic rate (Table 2). Our data also show, for gymnosperms, eudicots and monocots that there has also been an increase in TPU efficiency through evolutionary time (Table 1). However, there are too few data to compare other parameter values. It would be valuable to have data to compare parameters such as the half saturation constants to know if there have been evolutionary innovations in RuBisCO affinities. We restricted our search to in vivo parameter estimates. However, analysis of *K*_*c*_and *K*_*0*_ estimated using extracted enzyme in vitro has recently been done for many diverse lineages ranging from Bacteria, Archaea and Eukarea (Iñiguez *et al*., 2020). Interestingly, they find significant phylogenetic conservation with early groups such as Chorophyta (green algae), Cyanobacteria, and Euglenophyta (a group of photosynthetic flagellate algae) having RuBisCO affinities for CO_2_ that were very similar to vascular land plants, though the actual affinities ranged over an order of magnitude for these three groups of plants (Iñiguez *et al*., 2020). Thus, the global understanding of photosynthesis would benefit from more in vivo estimates of enzyme affinities.

Indeed, in terms of actual parameter estimates, we note that *C*_i_ (n=112), *Γ\** (n=64), *K*_*c*_ (n=43), *K*_*0*_ (n=29), and all parameters from eqns 4-5 were rarely estimated (Table 1,2, Fig S1, S2). Most studies use previously published estimates of these parameters to remove the degrees of freedom problem caused by over-parameterization of the FvCB model, and then simply estimate *V*_*cmax*_ (n=1294), *J*_*max*_ (n=891), and increasingly *T*_*p*_ (n=171). Parameters other than *V*_*cmax*_ and *J*_*cmax*_ are more challenging to estimate because of FvCB model over-parameterization. However, these more rarely estimated parameters are far from constant (Fig S1, S2) despite being required to solve the model. Thus, there is likely more variation in these more rarely estimated parameters (I.e *Γ\** n=64; *K*_*c*_, n=43; *K*_*0*_, n=29) than is currently known, and we suggest that more attention to these parameter estimates would greatly improve our ability to accurately model photosynthesis. Indeed, all parameters show either a skewed distribution with high kurtosis, or perhaps even a multi-modal distribution (Fig S1, S2). Currently, there are too few data to know whether these multi-modal distributions are artefacts of low sample sizes, or if they represent important physiological trade-offs.

It should also be noted that though we recorded respiration rates (*R*_d_, n=468) that were reported along with the main parameters of the FvCB equations, we did not seek out studies that report respiration rates of leaves. We are well aware that there are many more estimates of leaf respiration rates in the literature which are not associated with modelling exercises. However, like various resistances to CO_2_ movement into and through the leaf, seeking out leaf respiration rate measurements would require its own dedicated meta-analysis, which we view as a separate study. In the future, we seek to continue to update and expand this database of parameters. Future updates could also include accounting for stomatal and mesophyll conductance (Flexas *et al*., 2014, Niinemets *et al*., 2011).

In addition to the four orders of magnitude variation among species, the data also show that variation in parameter estimates within a species can be over an order of magnitude (Fig S3). Importantly, however, most of this variation can be explained by methodological differences among research groups and are driven by the leaf temperature and VPD inside the cuvette, as well as the nitrogen content of the leaves (i.e. the growing conditions of the plant). Variation among species is also partially explained by methods (Fig S4, S5) and clade (Fig 1,2, Table 1). This is well known, and not particularly surprising, so we do not dwell on these methodological effects here.

### Implications for global models

Most global models rely on simplifying the diversity of plant life by representing plants as some small number of average PFTs (Wullschleger *et al*., 2014). However, it is interesting thing to note that the average plant in a clade (Fig 2A, B), or PFT (Fig 2C, D) tells a different story than the diversity of results one sees when we model across species, genotypes and even individual leaves (Fig 2E, F). For example, from our modelled *A − C*_i_ curves, we can predict the average functional response of each major plant clade (Fig 1A) or PFT (Fig 1C) to rising CO_2_ levels. Because our data go back to the 1970s, the average partial pressure of intracellular CO_2_ across our data set was only 28.4 Pa (∼280 ppm), while at the time of this writing the partial pressure of CO_2_ in the atmosphere was approximately 42 Pa (∼415 ppm). The model output predicts that the average angiosperm has been able to take advantage of the increased CO_2_ in the atmosphere in this period from the 1970s to the present (Fig 2A). Furthermore, the PFT data show that the average angiosperm response is dominated largely by the average graminoids and average forbs, while the average tree and shrub shrubs were already at their maximum photosynthetic rate with CO_2_ at 28.4 Pa. If we continue this trend and cast forward to the IPCC predictions for the year 2100 (Ipcc, 2012) of 74 – 103.4 Pa (∼730-1020 ppm) CO_2_, our model results suggest that our hypothetical average plant representing different clades and PFTs are already near or at their maximum photosynthetic rates at CO_2_ levels of 42 Pa (Fig 1A, C). Our data using hypothetical mean PFTs suggest that only graminoids, have much capacity for increased CO_2_ assimilation beyond their current assimilation rates as we move toward the expected CO_2_ composition of the atmosphere by the year 2100. This seems to contradict empirical work where any plant species nearly always increase photosynthesis rates with increasing CO_2_ well beyond 42 Pa (Ainsworth & Long, 2005, Norby & Zak, 2011). What is the cause of this apparent contradiction?

We suggest, the lack of phylogenetic signal among all the species and clades (Fig 1A) means modelling plants as some hypothetical average member of a clade or PFT obscures important underlying species level variability. Indeed, when we examine the individual *A − C*_i_ curves for every leaf for which we had data, the model results show that almost every individual leaf has the capacity to dramatically increase its photosynthetic rate as we move from current CO_2_ levels to the IPCC predictions for the year 2100 (Fig 2E). Thus, our results tell two divergent stories: some hypothetical average plant from a clade (Fig 2A) or PFT (Fig 2C) is predicted to have little room for additional photosynthesis with increasing CO_2_, while actual empirically observed individual leaves almost all have significant room for additional photosynthesis with increasing CO_2_ (Fig 2E). Thus, the simplifying use of PFTs in most climate models may be dramatically underestimating the future photosynthetic capacity of terrestrial plants because using a mean PFT obscures the fact that most variation occurs within groups not among groups (Fig 1A). Ecologically, we would expect those species who can take advantage of increasing CO_2_ to expand in abundance resulting in an increased photosynthetic capacity at the community and ecosystem scale. The clade and PFT means do not capture this ecological change.

### Future directions: filling in gaps

Given the divergent results between the response of some mean hypothetical clade or PFT member which were almost at their maximum rate of photosynthesis in response to rising CO_2_, and the individual leaves which almost all had capacity to increase their photosynthetic rate in response to rising CO_2_, this seems like a problem for global models. On one hand, the use of a handful of PFTs is a necessary simplifying assumption in the face of a world with hundreds of thousands of species (Wullschleger *et al*., 2014). But on the other, it obscures the diversity of responses within each group which may drive future plant-climate feedbacks (Fig 1, 2). Newer models that include ecosystem demography and ecological competition among more types of plants are already likely the solution to this problem (Medvigy *et al*., 2009). We look forward to the increasing use and development of these ecosystem demography models which are a promising solution to this problem.

There are some holes in the global data that limit some conclusions that can be drawn. From a phylogenetic and biogeographic perspective, it seems that the lack of bryophyte and lichen parameter data (Fig 1A), particularly in the arctic (Fig 1B), is a hole in our ability to predict global photosynthesis. In some arctic and boreal systems, bryophytes and lichens can represent the bulk of net primary production by C_3_ pathways (Limpens *et al*., 2011). Gago *et al*. (2019) summarized empirically estimated values of *A*_*max*_, and show that mosses and liverworts fit into the evolutionary hierarchy such that their photosynthetic rates are lower than fern allies like lycophytes. The absence of FvCB parameters for bryophytes and lichens however, means that it is difficult to build the important results of Gago *et al*. (2019) into climate models. Since, most variability in the data occured within groups not among groups (Fig 1A), models would likely benefit from increased taxonomic coverage of these more rarely studied groups.

From a perspective of potential geographic bias, we cannot help but notice that – like many global datasets (E.g. Díaz *et al*., 2016, Luyssaert *et al*., 2007, Wright *et al*., 2004) – the FvCB parameter data are very western-centric, primarily coming from North American, Australian, and European species. This is likely a function of past funding levels, but we should work to correct this historical pattern. Indeed, many fields in science are well known to have a problem with diversity among members of the field (Swartz *et al*., 2019). Such lack of diversity is thought to limit perspective and cost the field bright minds from underrepresented minorities. Indeed, more diverse collaborations have been shown to lead to higher impact research (Alshebli *et al*., 2018). However, in a field like biology where diversity of life is also a part of the structure of our data, we suggest this western-centric data bias is also harming our global understanding of ecological systems as much as it is harming scientists from under-represented groups. There could be much to be gained by increased coverage of species from the South American, African, and Asian continents. Particularly, since most terrestrial biodiversity hotspots are in these regions. Data exists for almost every species of tree in the boreal forest, and arguably the majority of the most common trees in temperate forests, yet we know comparatively little about the diversity of photosynthetic responses of the enormous diversity of plant species in the tropics (Fig 1B). However, we do not think the solution to this dual problem with diversity of scientists and diversity of plant data is for western scientists to move their research into regions of Africa, South America and Asia that are under sampled. These regions already have scientists who can become experts or perhaps collaborators. Given that the cost of instruments required to estimate photosynthesis parameters is unusually high for ecological research, we recommend an international collaborative approach might be the most useful way to combine western access to expensive instruments, with local expertise in flora. Some have called such international collaborations the “fourth age of science” (Adams, 2013).

## Conclusion

Photosynthesis is a key process in the global climate system and is often modelled with FvCB type models. However, considering that there are 300,000 estimated plant species on earth, there is the potential for a large diversity of photosynthetic ability among plants. Thus, accurate models require large databases of parameter estimates. We have assembled such a database containing all parameter estimates required to solve the FvCB photosynthesis model. The publicly available database contains 3663 rows of data where at least one parameter was estimated for a given species. The summary includes parameter estimates from 359 different plant species from 96 taxonomic families representing all major vascular plant clades. The biogeographic coverage of species spans the majority of the globe, although there are some important gaps. We find very different predicted photosynthetic rates depending on whether we examine some hypothetical average plant that is meant to represent a clade or PFT, compared to when we model individual leaves in our dataset. Specifically, when hypothetical average plants are modelled we found that most clades are approaching their maximum photosynthetic rate in response to elevated CO_2_. However, when the breadth of responses for individual leaves are modelled, we found that almost all plants are predicted to increase photosynthetic rates in response to elevated CO_2_. We hope that this database can improve our understanding of global carbon flux through the terrestrial biosphere.

## ACKNOWLEDGMENTS

We thank Mike Mickelbart and Scott McAdam for many helpful discussions about photosynthesis, and we thank Laura Jessup and Abdel Halloway for comments on the manuscript. We thank Jaum Flexas and Joesteph Stinziano for detailed comments on the preprint of this manuscript. This work was funded in part by USDA NIFA Hatch funds to GGM (project number 1010722). The authors declare no conflicts of interest.

## DATA ACCESSABILITY

All data will be made publicly available on dryad upon publication at https://doi.org/10.5061/dryad.3tx95x6dr. All code used to make all figures and any data analysis are publicly available on GitHub (https://github.com/ggmcnickle/GlobalFvCB).

## AUTHOR CONTRIBUTIONS

Both authors contributed equally to data collection, analysis and writing.

