## Supplementary information for "A global database of photosynthesis model parameters, and modelled photosynthetic responses from every major terrestrial plant clade"

**SUPPLIMENTARY MATERIAL:**

Here we describe the search criteria and methods in the meta analysis. This section also includes more detailed supplementary tables and figures associated with the main text. This includes more detailed data summaries that at the taxonomic family level parameter (Table S1, S2), and by growth habit of each PFT (Table S3, and S4). We also show the raw frequency distribution of different parameter values (Fig S1, S2). Finally, we explore the source of variation both within and among species in parameter estimates. We find that large amounts of variation both within and among species can largely be explained by phylogeny, experimental methods and plant growing conditions (Table S5, Fig S3, S4).

**SUPPLIMENTARY METHODS**

*Literature search*

Parameter values for Eqns 1-6 were sought from empirical studies in the literature. In October 2017, we searched the Web of Science using the following “Farqhuar photosynthesis model” in the title only. This search returned 220 papers. We constrained the search to the Web of Science topics plant science, which reduced it to 162 papers. Each of these 162 papers were then critically read by author MR to remove unrelated papers captured by the search and identify those that contained parameter estimates. MR recorded experimental details, study species, basic experimental methods, and extracted parameter values for eqns 1-3. Subsequently, author GGM took a second pass through each of the 162 papers to check for any unusual or unique formulations of the FvCB model that could be expressed as Eqns 1-6. In addition, GGM used the bibliography of these papers to identify potential studies missing from the initial search. This identified 140 potential additional papers with parameter estimates. In turn, the bibliographies of this second set of papers were also inspected. This identified only one new study not already in our first or second search. The final set of data sources included parameters estimated from 147 unique references (Appendix A - Data sources).

One study (Kattge & Knorr, 2007) referenced the TRY database (Kattge *et al.*, 2020). So we made a data request for only publicly available data for $V_{c,max}$, $J_{max}$, $\Gamma^{*}$, and $K_{o}$ in June 2020. TRY returned data for 153 species, but inspection of the references associated with the TRY data entries revealed that we had already extracted data from the original reference for all but six of these entries. To avoid duplicates, we only recorded these six novel records. These six records have TRY as the reference, and the associated reference in the note column of the supplementary data file.

Finally, our search did not find any ferns or lycophytes. We hypothesized this might just be due to a difference in language across fields resulting in those data not appearing in our search. Indeed, through personal communication with a fern expert (S. McAdam, Pers. Comm.), we added literature containing parameter estimates for 33 fern and 2 lycophyte species. The addition of these 35 taxa were the only deviation from the systematic search described above. It is unlikely that we found 100% of parameter estimates available, but we view this as version 1.0 of a database of parameters that will be updated and maintained over time.

*Parameter value collection*

To collect parameter values from the pool of literature returned from the search and our fern expert, studies clearly must have estimated parameters from eqns 1-6 *in vivo*. For example, many studies estimate variables such as ‘quantum yield’ or ‘quantum efficiency’ or ‘light absorbance’ which are all terms occasionally associated with different parameters of eqns 4-5. However, to avoid confusion, we only collected data for the parameters of eqns 4-5 when it was clear that the parameter was estimated by actually fitting one of those three equations to data. In addition, we sought clear species level parameter estimates. Some studies estimated ecosystem or plant functional type (PFT) level estimates that were not species specific, or reported only the genus of the plant under study. If values for individual species could be identified in the paper, these were included; otherwise pooled ecosystem (PFT), or genus only estimates were excluded. In addition, we excluded studies that sought estimates of pure RuBisCO enzyme *in vitro*. We direct readers to Iñiguez *et al.* (2020) for a synthesis and meta-analysis of *in vitro* RuBisCO kinetic parameters.

Data were extracted most often from the text, tables, and figures. Many older studies simply state the parameter values in the text. A few older studies (n=3) reported parameters in the text only as ranges, and for these we recorded the midpoint of the range which represents the average. Data presented in tables were recorded as written. Data presented in figures were extracted with WebPlotDigitizer by calibrating the measurement tool to the $x$- and $y$-axis. Data in multiple panels with a common axis (e.g. $V_{c,max}$ vs Temperature and $J_{max}$ vs Temperature but in separate panels) were aligned on the common axis. For data that were reported in figures and aligned in this way, there were occasionally missing data points because of poor figure quality (E.g. too many over-lapping points that couldn’t be distinguished). Such missing data points are reported as NA in the data.

Many studies were experimental with any number of diverse treatments. For studies that included a clearly identifiable or labelled control (E.g. ambient temperature or CO_2_ vs elevated), only the control value was recorded. When there was no obvious control treatment (e.g. high and low nitrogen; during flowering and after flowering; stand age), we averaged groups. There are too many individual differences in treatments among hundreds of studies to describe all such decisions about treatment averaging here, but each decision was recorded in the note column of the supplementary data file along with the reference, and the location we collected data from within the reference (e.g. Fig $x$ or Table $y$).

In addition to photosynthesis parameters, leaf temperature and leaf nitrogen, we also recorded leaf mass per area (LMA) or specific leaf area (SLA) when reported as these too may influence leaf physiology. Finally, we also recorded the environmental conditions of the gas exchange cuvette which might have affected the parameter estimates. This included the irradiance used to fit $A-C_{i}$ curves, the partial pressure of CO_2_ used to fit $A-Q$ curves, and the relative humidity or vapour pressure deficit (VPD) of the cuvette if reported. As above, when a range was given, we took the midpoint as an average value. VPD was often reported as “VPD was kept below $x$” and in these cases we recorded $x$ as VPD. If it was stated that ‘ambient’ CO_2_ was used, but a value was not provided, the CO_2_ concentration in the atmosphere at the USA National Oceanic and Atmospheric Administration's Mauna Loa Observatory on Hawai'i was looked up for the summer of the publication year of the paper, otherwise we recorded ‘*NA*’. It should be noted that in many cases a value is not expected and those cells are blank in the supplementary data file. For example, a study which only estimated one of $V_{c,max}$ would only have measured an $A/C_{i}$ curve, and thus there would be no CO_2_ partial pressure for an $A/Q$ curve, no value of $J_{max}$, and so on.

*Treatment of pre-existing meta-analyses*

A few studies used the same dataset to explore new methods for parameter estimates. For example, Dreyer *et al.* (2001) took extremely detailed gas exchange measurements of seven species and estimated some FvCB parameters with those data. The authors freely shared their data, and a few subsequent studies explored more sophisticated parameter estimate methods on the same data (e.g. Kattge & Knorr, 2007; Su *et al.*, 2009). This was rare enough (n=3) that we decided to include all parameter estimates even though they emerged from the same data. The synthesis of data we present here includes such diverse methods spanning several decades of gas exchange and computing technology. We felt that the inclusion of parameters from a few studies that re-analysed old data, as well as the original estimates from those data, were representative of this diversity in methods. This is noted in the data file, so future users may exclude such repeat studies as they see fit for subsequent analyses.

*Standardisation of units*

Studies differed in the units reported for $C_{i}$, $\Gamma^{*}$, $K_{c}$, and $K_{o}$ which are either in units of atmospheric partial pressure or as CO_2_ concentration. To summarise the parameter estimates, we converted all parameters to a standard unit. Most commonly, these parameters were reported in units of partial pressure, and so we converted all estimates to Pa. Sometimes, these parameters were reported in units of concentration (e.g. μmol mol^-1^ or ppm). For the purposes of summary statistics, concentrations were converted into pressure using Dalton’s law of partial pressures assuming atmospheric pressure at sea level of 101.325 kPa such that 1 μmol mol^-1^ (or ppm) is equal to 0.101325 Pa. These are approximations and all original values and units are also given in the dataset in a separate column. Temperatures were converted to °C. Leaf nitrogen was converted to percent by dry weight (mg N g^-1^ DW) or to g m^-2^. Some studies report mean SLA while others report mean LMA. In the past, some authors simply convert mean SLA to mean LMA (or vice versa) by taking the inverse (E.g. Walker *et al.*, 2014; Díaz *et al.*, 2016). However, importantly the mean of $\left( x_{1},x_{2},\ldots,x_{n} \right)$ is not equivalent to the mean of $\left( \frac{1}{x_{1}},\frac{1}{x_{2}}, \ldots,\frac{1}{x_{n}} \right)$ by way of Jensen’s inequality (Jensen, 1906). Thus, to avoid introducing unnecessary error into our data we recorded SLA and LMA separately and do not attempt to interconvert them. We do however standardise their units to either m^2^ g^-1^ or g m^-2^ respectively. (Unlike converting mean SLA to LMA, unit conversions are mathematically acceptable because the mean of $c\left( x_{1},x_{2},\ldots,x_{n} \right)$ is equal to the mean of $\left( {cx}_{1},cx_{2},\ldots,{cx}_{n} \right)$, where $c$ is any constant used to convert units.) Finally, some older studies reported parameters in units of energy per leaf area per time (E.g. $Joules {cm}^{-2} s^{-1}$). This was also rare (n=2), and it was unclear how to adjust the units for comparison to the vast number of studies that express units of amount of CO_2_ per leaf area per time. These studies were not included.

*Variation within species*

To examine sources of variation in parameter estimates within a species, and its potential cause, we sought species for which at least 10 independent research groups had made measurements on the same species for either $V_{c,max}$ or $J_{max}$. We then used linear mixed effects models to investigate whether variable conditions in the cuvette, as well as biological properties of leaves explained the differences in parameter estimates within a species but across studies. In these models, species was treated as a random intercept to account for the effects of phylogeny.

*Variation among species*

To examine sources of variation among species we used Principle Components Analysis (PCA) on the pairwise complete correlation matrix to examine correlations among variables in our dataset.

**SUPPLIMENTARY RESULTS**

*Variation within species*

There were five species for which 10 or more studies had independently estimated either $V_{c,max}$ or $J_{max}$. The data show that estimates of these two parameters within a species can vary over an order of magnitude (Fig S3A, B).

Though we had LMA, SLA, light levels used for saturating light measurements, and CO2 pressures used for saturating carbon measurements for many species, we did not have enough data for the five species in this part of the analysis to include these variables in the regression. The mixed effets multiple regression therefore only included VPD, leaf N (per area), and leaf temperature. Unfortunately, for these five species, most had been measured at only two temperatures, which meant that temperature was not significant. Therefore, multiple regression revealed that for $V_{c,max}$ and $J_{max}$ the variation within a species for these five species was driven by the nitrogen content of the leaves and the VPD tolerance of the measurements (Table S5). Therefore, even though estimates within a species can vary over an order of magnitude: we conclude that this variation can be explained largely by methodological differences among research groups, and growing conditions of the plants under study that affect leaf nitrogen.

*Variation among species*

Among the different variables we collected, only temperature, VPD, Leaf N per area, and per mass, saturating light level for the A-C curve (Q), and saturating CO_2_ level for the A-Q curve had enough correlated observations to estimate a pairwise correlation matrix (Fig S4). Similarly, among the different parameters, only $V_{c,max}$, $J_{max}$, $C_{i}$, $\Gamma^{*}$, $R_{d}$, and $T_{p}$ had enough linked observations to estimate a correlation matrix. Using PCA, the first two PC axes explained 68.5% of the variation among these variables and parameters (Fig 4). The PCA shows the well-known positive correlation between temperature $V_{c,max}$, $J_{max}$ and $T_{p}$ which has been studied many times (Fig S5). However, it also revealed a positive relationship between $C_{i}$ and $\Gamma^{*}$, and that both of these were negatively correlated with leaf nitrogen content and the CO_2_ level used to fit the A-Q curve (Fig S4).

**TABLE S1**: Taxonomic summary of FvCB parameter values across clade, and expanded across family for Eqns 1-3, 6 and 7. The mean is given if available, blank cells represent no data. The standard deviation, and sample size are shown in parentheses after the mean. That is, values in each cell represent: mean (SD, n). Blank cells mean no data is available. For sample sizes less than three, the standard deviation is reported as NA.

| **Clade** | **Family** | $\boldsymbol{V}_{\boldsymbol{c max}}$  **(μmol m^-2^ s^-1^)** | $\boldsymbol{J}_{\boldsymbol{max}}$  **(μmol m^-2^ s^-1^)** | $\boldsymbol{C}_{\boldsymbol{i}}$  **(Pa)** | $\boldsymbol{\Gamma}^{\boldsymbol{*}}$  **(Pa)** | $\boldsymbol{K}_{\boldsymbol{c}}$  **(Pa)** | $\boldsymbol{K}_{\boldsymbol{o}}$  **(Pa)** | $\boldsymbol{R}_{\boldsymbol{d}}$  **(μmol m^-2^ s^-1^)** | $\boldsymbol{T}_{\boldsymbol{p}}$  **(μmol m^-2^ s^-1^)** |
| --- | --- | --- | --- | --- | --- | --- | --- | --- | --- |
| **Lycophyte** | Lycopodiaceae | 17.15 (NA, 2) | 25.5 (NA, 2) |  |  |  |  | 0.5 (NA, 2) | |
| **Fern** | **ALL** | **51.2 (27.9, 33)** | **59.7 (24.2, 33)** |  |  |  |  | **0.5 (0.3, 33)** | |
|  | Aspleniaceae | 65 (NA, 2) | 65.6 (NA, 2) |  |  |  |  | 1 (NA, 2) | |
|  | Athyriaceae | 55 (NA, 1) | 70.2 (NA, 1) |  |  |  |  | 1 (NA, 1) | |
|  | Blechnaceae | 37.2 (6.8, 4) | 50.9 (9.9, 4) |  |  |  |  | 0.1 (0, 4) | |
|  | Cystopteridaceae | 34.5 (NA, 2) | 52.6 (NA, 2) |  |  |  |  | 0.5 (NA, 2) | |
|  | Dennstaedtiaceae | 65.7 (NA, 2) | 88.3 (NA, 2) |  |  |  |  | 0.5 (NA, 2) | |
|  | Dicksoniaceae | 38.8 (NA, 1) | 60.3 (NA, 1) |  |  |  |  | 0.4 (NA, 1) | |
|  | Dryopteridaceae | 58.7 (14.9, 5) | 67.5 (6.5, 5) |  |  |  |  | 0.4 (0.4, 5) | |
|  | Equisetaceae | 54.1 (10.4, 4) | 60.7 (23, 4) |  |  |  |  | 0.9 (0.2, 4) | |
|  | Gleicheniaceae | 28.5 (NA, 1) | 53.7 (NA, 1) |  |  |  |  | 0.3 (NA, 1) | |
|  | Lygodiaceae | 19.8 (NA, 1) | 19.8 (NA, 1) |  |  |  |  | 0.2 (NA, 1) | |
|  | Nephrolepidaceae | 17.4 (NA, 1) | 22.2 (NA, 1) |  |  |  |  | 0.2 (NA, 1) | |
|  | Onocleaceae | 35.8 (NA, 1) | 51 (NA, 1) |  |  |  |  | 0.6 (NA, 1) | |
|  | Ophioglossaceae | 19.5 (NA, 2) | 22.5 (NA, 2) |  |  |  |  | 0.6 (NA, 2) | |
|  | Osmundaceae | 69.8 (NA, 1) | 69.7 (NA, 1) |  |  |  |  | 0.2 (NA, 1) | |
|  | Pteridaceae | 98.7 (62.3, 3) | 86.8 (50.4, 3) |  |  |  |  | 0.4 (0.2, 3) | |
|  | Thelypteridaceae | 50.1 (NA, 2) | 61.4 (NA, 2) |  |  |  |  | 0.5 (NA, 2) | |
| **Gymnosperm** | **ALL** | **34.7 (32.25, 461)** | **65.2 (42.93, 301)** | **19.4 (5.14, 37)** | **7.7 (3.04, 27)** | **27.4 (NA, 2)** | **41543.3 (NA, 2)** | **0.9 (0.45, 41)** | **2.8 (2.62, 49)** |
|  | Cupressaceae | 24.6 (NA, 1) |  |  |  |  |  |  |  |
|  | Pinaceae | 34.7 (32.26, 457) | 65.3 (43.08, 298) | 19.4 (5.14, 37) | 7.7 (3.04, 27) | 27.4 (NA, 2) | 41543.3 (NA, 2) | 0.9 (0.45, 41) | 2.8 (2.62, 49) |
|  | Podocarpaceae | 44.8 (40.53, 3) | 59.4 (28.06, 3) |  |  |  |  |  |  |
| **Magnoliid** | **ALL** | **21 (12.85, 18)** | **119.3 (29.16, 3)** |  |  |  |  | **0.7 (0, 1)** |  |
|  | Lauraceae | 15.6 (5.73, 15) |  |  |  |  |  |  |  |
|  | Myristicaceae | 51 (NA, 1) | 153 (NA, 1) |  |  |  |  | 0.7 (NA, 1) |  |
|  | Piperaceae | 41 (NA, 2) | 102.5 (NA, 2) |  |  |  |  |  |  |
| **Eudicot** | **ALL** | **76.3 (56.88, 2037)** | **135.9 (64.8, 1060)** | **28.8 (5.67, 43)** | **4.3 (1.87, 63)** | **47.5 (31.63, 42)** | **26442.6 (11382.05, 31)** | **1.5 (1.76, 283)** | **6.8 (2.6, 48)** |
|  | Acanthaceae | 85 (NA, 1) | 145 (NA, 1) |  |  |  |  |  |  |
|  | Altingiaceae | 52.4 (29.56, 145) | 144.6 (31.32, 11) |  |  |  |  |  |  |
|  | Amaranthaceae | 112.2 (60.7, 18) | 223.6 (76.45, 7) | 36.8 (NA, 1) | 3.8 (0.92, 4) | 46 (NA, 1) |  | 0.6 (0.73, 6) |  |
|  | Anacardiaceae | 77 (NA, 2) | 144 (NA, 2) |  |  |  |  |  |  |
|  | Apiaceae |  |  | 35.7 (NA, 1) |  |  |  |  |  |
|  | Apocynaceae | 82.5 (NA, 2) | 157 (NA, 2) |  |  |  |  |  |  |
|  | Aquifoliaceae | 9.5 (NA, 1) |  |  |  |  |  |  |  |
|  | Araliaceae | 36 (NA, 1) | 69 (NA, 1) |  |  |  |  |  |  |
|  | Asteraceae | 90.8 (62.26, 28) | 156.7 (78.06, 19) | 32.1 (5.67, 7) | 0.6 (NA, 2) | 28.7 (NA, 1) |  | 3.8 (4.8, 7) |  |
|  | Betulaceae | 62.2 (46.24, 80) | 115.9 (70.69, 82) | 25.7 (2.38, 7) | 4 (NA, 1) | 49.6 (NA, 1) | 28620 (NA, 1) | 1.6 (1.35, 6) | 6.6 (1.6, 6) |
|  | Bignoniaceae | 45 (NA, 1) | 71 (NA, 1) |  |  |  |  |  |  |
|  | Brassicaceae | 86.5 (43.23, 18) | 154.3 (51.56, 12) |  | 4.5 (0.92, 7) | 26.5 (NA, 1) | 20100 (NA, 1) | 0.4 (NA, 1) | 6.5 (NA, 2) |
|  | Buxaceae | 27 (NA, 1) | 69 (NA, 1) |  |  |  |  |  |  |
|  | Campanulaceae | 56.8 (NA, 2) | 80 (NA, 1) |  |  |  |  |  |  |
|  | Cercidiphyllaceae | 32.3 (20.47, 105) |  |  |  |  |  |  |  |
|  | Cistaceae | 71 (NA, 1) | 167 (NA, 1) |  |  |  |  |  |  |
|  | Combretaceae | 68 (11, 3) | 129 (74.22, 3) |  |  |  |  | 1 (0.21, 3) |  |
|  | Convolvulaceae | 26 (NA, 1) |  |  |  |  |  |  |  |
|  | Cucurbitaceae | 32.7 (16, 3) | 118.7 (NA, 2) |  |  |  |  | 0.3 (NA, 1) | 12.3 (NA, 1) |
|  | Cunoniaceae | 29 (NA, 2) | 39.9 (NA, 2) |  |  |  |  |  |  |
| **Eudicot (continued)** | Daphniphyllaceae | 16.2 (NA, 1) | 35.1 (NA, 1) |  |  |  |  |  |  |
|  | Dipterocarpaceae | 30.3 (11.85, 13) | 63.8 (32.18, 13) |  |  |  |  |  |  |
|  | Elaeagnaceae | 52.3 (NA, 1) | 111.8 (NA, 1) |  | <0.1 (NA, 1) |  |  | 4.2 (NA, 1) |  |
|  | Elaeocarpaceae | 31.6 (15.99, 30) | 71.6 (33.26, 30) |  |  |  |  |  |  |
|  | Ericaceae | 26.8 (23.49, 28) | 86.6 (25.3, 5) |  |  | 50.1 (36.25, 6) | 33388.7 (14890.7, 6) | 0.8 (0.24, 6) |  |
|  | Euphorbiaceae | 45 (36.18, 4) | 97.8 (74.03, 4) |  |  |  |  |  |  |
|  | Fabaceae | 81.6 (36.95, 100) | 181.5 (69.49, 53) | 24.3 (NA, 2) | 5.1 (2.11, 8) | 43.9 (24.17, 9) | 25075.9 (12920.11, 9) | 1.9 (1.64, 17) | 11.3 (NA, 2) |
|  | Fagaceae | 52.1 (36.08, 295) | 107 (57.94, 175) |  | 3.7 (NA, 2) | 60.8 (NA, 2) |  | 1.9 (2.95, 50) | 5.3 (1.49, 7) |
|  | Garryaceae | 9 (NA, 1) |  |  |  |  |  |  |  |
|  | Geraniaceae | 52.4 (NA, 1) |  |  |  |  |  |  |  |
|  | Goupiaceae | 27 (NA, 1) | 68 (NA, 1) |  |  |  |  |  |  |
|  | Hypericaceae |  |  | 36.8 (NA, 1) |  |  |  |  |  |
|  | Juglandaceae | 57.5 (44.21, 37) | 99.2 (45.8, 37) |  | 3.7 (NA, 1) | 61.1 (NA, 1) |  | 0.9 (0.78, 26) | 4.5 (NA, 1) |
|  | Lamiaceae |  |  | 31.2 (NA, 1) |  |  |  |  |  |
|  | Lauraceae | 27.6 (13.49, 8) |  |  |  |  |  |  |  |
|  | Lythraceae |  |  | 35.7 (NA, 1) |  |  |  |  |  |
|  | Malvaceae | 99.1 (51.3, 46) | 164.3 (70.34, 40) |  |  |  |  | 1.4 (0.94, 14) | 12.1 (NA, 2) |
|  | Marantaceae | 124 (NA, 1) | 96 (NA, 1) |  |  |  |  | 1.3 (NA, 1) |  |
|  | Meliaceae | 19 (NA, 2) | 49 (NA, 2) |  |  |  |  |  |  |
|  | Moraceae | 33 (NA, 1) | 70 (NA, 1) |  |  |  |  |  |  |
|  | Myrtaceae | 56.3 (27.44, 78) | 122.8 (38.68, 71) | 28.6 (2.82, 12) | 3.9 (0.58, 3) | 30.6 (NA, 1) | 25939.2 (NA, 1) | 0.3 (0.61, 21) | 8.5 (NA, 1) |
|  | Nyssaceae | 17 (NA, 1) | 32 (NA, 1) |  |  |  |  |  |  |
|  | Oleaceae | 75.6 (46.01, 13) | 131.3 (58.84, 12) |  |  |  |  | 2 (0.9, 8) | 5.3 (NA, 1) |
|  | Onagraceae | 51.6 (24.77, 29) | 107.6 (40.44, 29) |  |  |  |  |  |  |
| **Eudicot (continued)** | Orobanchaceae | 52.3 (NA, 1) |  |  |  |  |  |  |  |
|  | Paracryphiaceae | 16.9 (NA, 1) | 50.7 (NA, 1) |  |  |  |  |  |  |
|  | Pentaphylacaceae | 4.6 (NA, 1) |  |  |  |  |  |  |  |
|  | Plantaginaceae | 53.6 (7.44, 4) | 200 (NA, 1) | 36.7 (NA, 1) |  |  |  |  |  |
|  | Platanaceae | 72.9 (36.17, 149) |  |  |  |  |  |  |  |
|  | Polygonaceae | 43.3 (5.19, 7) | 87 (8.54, 4) | 33.8 (2.7, 3) |  |  |  |  |  |
|  | Primulaceae | 71 (NA, 1) | 135 (NA, 1) |  |  |  |  |  |  |
|  | Proteaceae | 44.6 (38.59, 5) | 58 (15.08, 4) |  |  |  |  |  |  |
|  | Ranunculaceae | 43 (24.35, 4) | 85 (NA, 1) |  |  |  |  |  |  |
|  | Rhamnaceae | 42 (NA, 1) | 94 (NA, 1) |  |  |  |  |  |  |
|  | Rosaceae | 80.5 (55.1, 236) | 139.7 (54.9, 99) | 18 (NA, 1) | 9.1 (NA, 1) | 21.1 (NA, 1) |  | 0.9 (0.37, 42) | 6 (1.69, 20) |
|  | Rubiaceae | 31 (NA, 2) | 72.5 (NA, 2) |  |  |  |  |  |  |
|  | Rutaceae | 51.9 (46.27, 9) | 73.4 (43.5, 9) |  |  |  |  |  |  |
|  | Salicaceae | 145 (59.86, 364) | 188.4 (38.26, 214) | 29.7 (NA, 1) | 4.7 (1.72, 6) | 70.8 (33.71, 6) | 23448.8 (NA, 2) | 2.3 (1.18, 44) |  |
|  | Sapindaceae | 39 (42.15, 61) | 110.9 (52.21, 18) |  |  |  |  | 1.4 (0.98, 12) | 5.2 (NA, 2) |
|  | Scrophulariaceae | 91 (NA, 1) | 281 (NA, 1) |  |  |  |  |  |  |
|  | Simmondsiaceae | 32 (NA, 1) | 91 (NA, 1) |  |  |  |  |  |  |
|  | Solanaceae | 65.7 (64.09, 30) | 162.6 (79.26, 25) | 21.1 (NA, 2) | 4.5 (1.85, 26) | 41.2 (42.79, 11) | 24570.5 (9497.45, 11) | 1.2 (0.76, 13) | 10.5 (NA, 2) |
|  | Tamaricaceae | 12 (NA, 1) |  |  |  |  |  |  |  |
|  | Theaceae | 32 (NA, 1) | 71 (NA, 1) |  |  |  |  |  |  |
|  | Ulmaceae | 49 (NA, 1) | 89 (NA, 1) |  |  |  |  | 0.7 (NA, 1) |  |
|  | Urticaceae | 50 (NA, 1) | 133 (NA, 1) |  |  |  |  | 1 (NA, 1) |  |
|  | Vitaceae | 57.2 (22.99, 47) | 91.1 (33.9, 47) | 20 (NA, 1) | 4.1 (NA, 1) |  |  | 0.2 (NA, 1) | 5.9 (NA, 1) |
|  | Zygophyllaceae | 93.6 (NA, 2) | 159.9 (NA, 2) | 18.6 (NA, 1) |  | 30.5 (NA, 1) |  | 2.9 (NA, 1) |  |
| **Monocot** | **ALL** | **85.9 (41.06, 400)** | **197.4 (93.85, 352)** | **31.7 (6.99, 27)** | **1.9 (1.97, 8)** | **51.9 (32.59, 22)** | **37944 (17120.35, 20)** | **0.8 (0.52, 228)** | **13 (5.35, 236)** |
|  | Araceae | 59.8 (24.08, 5) | 94.9 (62.65, 5) | 23.2 (NA, 1) |  |  |  | 0.1 (NA, 1) |  |
|  | Asparagaceae | 88.5 (NA, 1) | 125.1 (NA, 1) | 20.1 (NA, 1) |  | 24.6 (NA, 1) |  | 1.9 (NA, 1) |  |
|  | Cyperaceae | 31 (NA, 1) |  | 33.6 (2.27, 5) |  |  |  |  |  |
|  | Iridaceae |  |  | 32.2 (NA, 1) |  |  |  |  |  |
|  | Marantaceae | 52 (NA, 1) | 78 (NA, 1) |  |  |  |  | 0.5 (NA, 1) |  |
|  | Poaceae | 87.2 (41.1, 388) | 199.6 (93.04, 342) | 26.9 (8.34, 7) | 2.96 (1.66, 5) | 53.6 (32.85, 21) | 37944 (17120.35, 20) | 0.8 (0.49, 222) | 13 (5.35, 236) |

**TABLE S2**: Taxonomic summary of FvCB parameter values across clade, and family, and family for Eqns 4-5. The mean is given if available. The standard deviation, and sample size are shown in parentheses after the mean. That is, values in each cell represent: mean (SD,n). Blank cells mean no data is available. This table includes all families shown in Table S1 for ease of comparison, however, the parameters of Eqns 4-5 are rarely estimated, so there are many empty rows. For sample sizes less than three, the standard deviation is reported as NA.

| **Clade** | **Family** | $\boldsymbol{\alpha}$ **(Eqn 4)** | $\boldsymbol{\theta}$ **(Eqn 4)** | $\boldsymbol{a}$ **(Eqn 5)** |
| --- | --- | --- | --- | --- |
| **Gymnosperm** | **ALL** | **0.23 (0.05, 14)** | **0.53 (0.17, 13)** | **0.19 (0.02, 3)** |
|  | Cupressaceae |  |  |  |
|  | Pinaceae | 0.23 (0.05, 14) | 0.53 (0.17, 13) | 0.19 (0.02, 3) |
|  | Podocarpaceae |  |  |  |
| **Magnoliid** | **ALL** |  | **0.06 (NA, 1)** |  |
|  | Piperaceae | |  |  |
|  | Lauraceae | |  |  |
|  | Myristicaceae | | 0.06 (NA, 1) |  |
| **Eudicot** | **ALL** | **0.27 (0.22, 26)** | **0.51 (0.41, 44)** | **0.16 (0.09, 3)** |
|  | Acanthaceae |  |  |  |
|  | Altingiaceae |  |  |  |
|  | Amaranthaceae | 0.02 (NA, 1) |  |  |
|  | Anacardiaceae |  |  |  |
|  | Apiaceae |  |  |  |
|  | Apocynaceae |  |  |  |
|  | Aquifoliaceae |  |  |  |
|  | Araliaceae |  |  |  |
|  | Asteraceae | 0.31 (NA, 2) | 0.41 (0.38, 4) |  |
|  | Betulaceae | 0.16 (NA, 1) | 0.92 (NA, 1) |  |
|  | Bignoniaceae |  |  |  |
|  | Brassicaceae |  |  |  |
|  | Buxaceae |  |  |  |
|  | Campanulaceae |  |  |  |
|  | Cercidiphyllaceae |  |  |  |
|  | Cistaceae |  |  |  |
|  | Combretaceae | 0.03 (0, 3) |  |  |
|  | Cucurbitaceae |  |  |  |
|  | Cunoniaceae |  |  |  |
|  | Daphniphyllaceae |  |  |  |
|  | Dipterocarpaceae |  |  |  |
|  | Elaeagnaceae |  |  |  |
|  | Elaeocarpaceae |  | 0.1 (NA, 1) |  |
|  | Ericaceae |  |  |  |
|  | Euphorbiaceae |  |  | 0.06 (NA, 1) |
|  | Fabaceae |  |  |  |
|  | Fagaceae | 0.08 (NA, 2) | 0.66 (0.42, 10) |  |
|  | Garryaceae | 0.06 (NA, 1) |  |  |
|  | Geraniaceae |  |  |  |
|  | Goupiaceae |  |  |  |
|  | Hypericaceae |  |  |  |
|  | Juglandaceae |  |  |  |
|  | Lamiaceae |  |  |  |
|  | Lauraceae |  |  |  |
|  | Lythraceae |  |  |  |
|  | Malvaceae |  |  |  |
|  | Marantaceae | 0.28 (NA, 1) | 0.63 (NA, 1) |  |
|  | Meliaceae |  | 0.09 (NA, 1) |  |
|  | Moraceae |  |  |  |
|  | Myrtaceae |  |  |  |
|  | Nyssaceae | 0.6 (NA, 2) | 0.16 (0.28, 10) |  |
|  | Oleaceae |  |  |  |
|  | Onagraceae |  |  |  |
|  | Orobanchaceae |  |  |  |
| **Eudicot** | Paracryphiaceae |  |  |  |
| **(continued)** | Pentaphylacaceae |  |  |  |
|  | Plantaginaceae |  |  |  |
|  | Platanaceae |  |  |  |
|  | Polygonaceae |  |  |  |
|  | Primulaceae |  |  |  |
|  | Proteaceae |  |  |  |
|  | Ranunculaceae |  |  |  |
|  | Rhamnaceae |  |  |  |
|  | Rosaceae |  |  |  |
|  | Rubiaceae |  | 0.1 (NA, 1) |  |
|  | Rutaceae |  |  |  |
|  | Salicaceae |  |  |  |
|  | Sapindaceae | 0.17 (NA, 1) |  | 0.23 (NA, 1) |
|  | Scrophulariaceae |  |  |  |
|  | Simmondsiaceae |  |  |  |
|  | Solanaceae |  |  |  |
|  | Tamaricaceae | 0.39 (0.11, 11) | 0.84 (0.2, 12) |  |
|  | Theaceae | 0.02 (NA, 1) |  |  |
|  | Ulmaceae |  |  |  |
|  | Urticaceae |  | 0.04 (NA, 1) |  |
|  | Vitaceae |  | 0.04 (NA, 1) |  |
|  | Zygophyllaceae |  |  | 0.18 (NA, 1) |
|  | Convolvulaceae |  |  |  |
| **Monocot** | **ALL** | **0.27 (NA, 2)** | **0.67 (0.18, 230)** |  |
|  | Araceae | 0.1 (NA, 1) | 0.91 (NA, 2) |  |
|  | Asparagaceae |  |  |  |
|  | Cyperaceae |  |  |  |
|  | Iridaceae |  |  |  |
|  | Juncaceae |  |  |  |
|  | Marantaceae |  | 0.06 (NA, 1) |  |
|  | Poaceae | 0.44 (NA, 1) | 0.67 (0.18, 227) |  |

Note: Lycophytes and Ferns are not shown because there were no data for these clades.

**TABLE S3**: Functional group summary of FvCB parameter values across type and growth habit for Eqns 1-3. The mean is given if available. The standard deviation, and sample size are shown in parentheses after the mean. That is, values in each cell represent: mean (SD,n). Blank cells mean no data is available. For sample sizes less than three, the standard deviation is reported as NA. For a fern functional type, see trachaeophytes in Table 2.

| **Type** | **Habit** | $\boldsymbol{V}_{\boldsymbol{c max}}$  **(μmol m^-2^ s^-1^)** | $\boldsymbol{J}_{\boldsymbol{max}}$  **(μmol m^-2^ s^-1^)** | $\boldsymbol{C}_{\boldsymbol{i}}$  **(Pa)** | $\boldsymbol{\Gamma}^{\boldsymbol{*}}$  **(Pa)** | $\boldsymbol{K}_{\boldsymbol{c}}$  **(Pa)** | $\boldsymbol{K}_{\boldsymbol{o}}$  **(Pa)** | $\boldsymbol{R}_{\boldsymbol{d}}$  **(μmol m^-2^ s^-1^)** | $\boldsymbol{T}_{\boldsymbol{p}}$  **(μmol m^-2^ s^-1^)** |
| --- | --- | --- | --- | --- | --- | --- | --- | --- | --- |
| **C3 Graminoid** | **ALL** | **87.2 (41.1, 388)** | **199.6 (92.2, 342)** | **29.7 (7.1, 12)** | **3 (1.7, 5)** | **53.6 (32.8, 21)** | **37944 (17120.3, 20)** | **0.8 (0.5, 222)** | **13 (5.4, 236)** |
|  | Annual | 78.4 (NA, 1) |  | 20.3 (6, 3) |  |  |  |  |  |
|  | Crop | 94.7 (39.4, 336) | 204.1 (89.7, 302) |  | 3.7 (0.2, 4) | 53.6 (32.8, 21) | 37944 (17120.3, 20) | 0.8 (0.5, 221) | 13 (5.4, 236) |
|  | Perennial | 50.1 (27.8, 51) | 174.3 (102.8, 40) | 32.9 (4, 9) | 0 (NA, 1) |  |  | 0.5 (NA, 1) |  |
| **Forb** | **ALL** | **83.5 (50.3, 222)** | **167.5 (75.4, 123)** | **31.2 (6, 22)** | **4.3 (1.6, 43)** | **41.1 (32.2, 23)** | **24574.4 (10685.8, 21)** | **1.5 (2.2, 40)** | **9.4 (3.4, 6)** |
|  | Annual | 128.7 (60.9, 25) | 187.9 (105.1, 12) | 34.1 (2, 7) | 4.5 (0.9, 7) | 26.5 (NA, 1) | 20100 (NA, 1) | 0.4 (NA, 1) | 6.5 (NA, 2) |
|  | Biennial | 129.4 (64.4, 3) | 225.7 (103.5, 3) |  |  |  |  | 0.7 (NA, 1) |  |
|  | Crop | 81.6 (47.6, 145) | 161.5 (62.3, 81) | 22.8 (3.9, 5) | 4.4 (1.7, 35) | 42.7 (33.4, 21) | 24809.9 (10925.1, 20) | 1.3 (1.2, 33) | 10.9 (3.2, 4) |
|  | Perennial | 62.6 (33.9, 47) | 173.6 (93, 26) | 33.3 (5, 10) | 0.6 (NA, 1) | 24.6 (NA, 1) |  | 3.6 (5.5, 5) |  |
| **Vine** | **ALL** | **55.9 (23.1, 55)** | **95.4 (37.2, 54)** | **20 (NA, 1)** | **4.1 (NA, 1)** |  |  | **0.5 (0.3, 5)** | **9.1 (NA, 2)** |
|  | Annual | 26 (NA, 1) |  |  |  |  |  |  |  |
|  | Crop | 56.2 (23.3, 51) | 92.7 (34.4, 50) | 20 (NA, 1) | 4.1 (NA, 1) |  |  | 0.2 (NA, 2) | 9.1 (NA, 2) |
|  | Perennial | 63.5 (16.3, 3) | 140.7 (60.2, 3) |  |  |  |  | 0.7 (0.1, 3) |  |
| **Shrub** | **ALL** | **72.1 (58.4, 166)** | **125.7 (63.3, 143)** | **18.7 (0.8, 3)** | **2.8 (3.5, 4)** | **26.8 (5, 3)** |  | **2.1 (1.7, 23)** | **6.6 (2.5, 21)** |
|  | Crop | 84.5 (41.8, 39) | 158.1 (61, 40) |  |  |  |  | 1.4 (1, 13) | 12.1 (NA, 2) |
|  | Deciduous | 76.4 (68.8, 93) | 116.2 (62.3, 78) | 18.8 (NA, 2) | 2.8 (3.5, 4) | 24.9 (NA, 2) |  | 3.2 (2.2, 8) | 6 (1.7, 19) |
|  | Evergreen | 46.8 (34.8, 34) | 102.7 (50.5, 25) | 18.6 (NA, 1) |  | 30.5 (NA, 1) |  | 2 (NA, 2) |  |
| **Tree** | **ALL** | **66.6 (56.1, 2077)** | **114.4 (65.4, 1048)** | **22.4 (6, 57)** | **6.6 (3, 42)** | **54.9 (30.1, 19)** | **32073 (12035.2, 12)** | **1.3 (1.6, 259)** | **3.7 (2.7, 68)** |
|  | Crop | 75.5 (50.1, 86) | 127.1 (54.4, 83) |  | 9.1 (NA, 1) |  |  | 0.9 (0.4, 40) | 6.1 (NA, 1) |
|  | Deciduous | 77.8 (59.7, 1424) | 139.8 (64.1, 588) | 26.2 (2.6, 8) | 4.7 (1.8, 11) | 65.7 (26.6, 10) | 25172.5 (7563, 3) | 1.8 (1.9, 150) | 5.7 (1.5, 17) |
|  | Evergreen | 37.5 (32.3, 561) | 74.3 (47, 372) | 21.7 (6.1, 49) | 7.3 (3.1, 30) | 42.9 (30.7, 9) | 34373.1 (12690.7, 9) | 0.7 (0.6, 69) | 2.9 (2.7, 50) |

Note: For a ‘fern’ or ‘lycophyte’ PFT, simply use the values reported in Table 2

**TABLE S4**: Functional group summary of FvCB parameter values across type, and growth habit for Eqns 4-5. The mean is given if available. The standard deviation, and sample size are shown in parentheses after the mean. That is, values in each cell represent: mean (SD, n). Blank cells mean no data is available. For sample sizes less than three, the standard deviation is reported as NA.

| **Type** | **Habit** | $\boldsymbol{\alpha}$ **(Eqn 4)** | $\boldsymbol{\theta}$ **(Eqn 4)** | $\boldsymbol{a}$ **(Eqn 5)** |
| --- | --- | --- | --- | --- |
| **C3 Graminoid** | **ALL** |  | **0.68 (0.2, 220)** |  |
|  | Annual |  |  |  |
|  | Crop |  | 0.69 (0.1, 217) |  |
|  | Perennial |  | 0.55 (0.4, 3) |  |
| **Forb** | **ALL** | **0.32 (0.2, 17)** | **0.77 (0.3, 23)** |  |
|  | Annual |  |  |  |
|  | Biennial |  |  |  |
|  | Crop | 0.32 (0.2, 15) | 0.82 (0.2, 17) |  |
|  | Perennial |  | 0.6 (0.5, 5) |  |
| **Vine** | **ALL** |  | **0.65 (0.5, 3)** |  |
|  | Annual |  |  |  |
|  | Crop |  |  |  |
|  | Perennial |  | 0.65 (0.5, 3) |  |
| **Shrub** | **ALL** | **0.02 (0, 4)** | **0.33 (0.5, 4)** |  |
|  | Crop |  |  |  |
|  | Deciduous |  | 0.33 (0.5, 4) |  |
|  | Evergreen |  |  |  |
| **Tree** | **ALL** | **0.24 (0.2, 20)** | **0.33 (0.3, 30)** | **0.17 (0.1, 5)** |
|  | Crop |  |  |  |
|  | Deciduous | 0.11 (0.1, 4) | 0.2 (0.4, 7) |  |
|  | Evergreen | 0.28 (0.2, 16) | 0.4 (0.3, 21) | 0.16 (0.1, 4) |
|  | NA |  |  |  |

Note: Lycophytes and Ferns are not shown because there were no data for these clades.

**TABLE S5**: ANOVA table for the GLMM on within species variation from a type III sum of squares, and Satterthwaite's method for estimating denominator degrees of freedom (df). Temperature was initially included, but was not significant in either model, and model fit was increased by dropping temperature.

| **Parameter** | **Factor** | **Num. df** | **Den. df** | **F** | **p** |
| --- | --- | --- | --- | --- | --- |
| **V** | **VPD** | 1 | 108.7 | 29 | <0.0001 |
|  | **Leaf N** | 1 | 108.2 | 26 | <0.0001 |
| **J** | **VPD** | 1 | 100 | 4.9 | 0.0286 |
|  | **Leaf N** | 1 | 96.6 | 13.2 | 0.0004 |

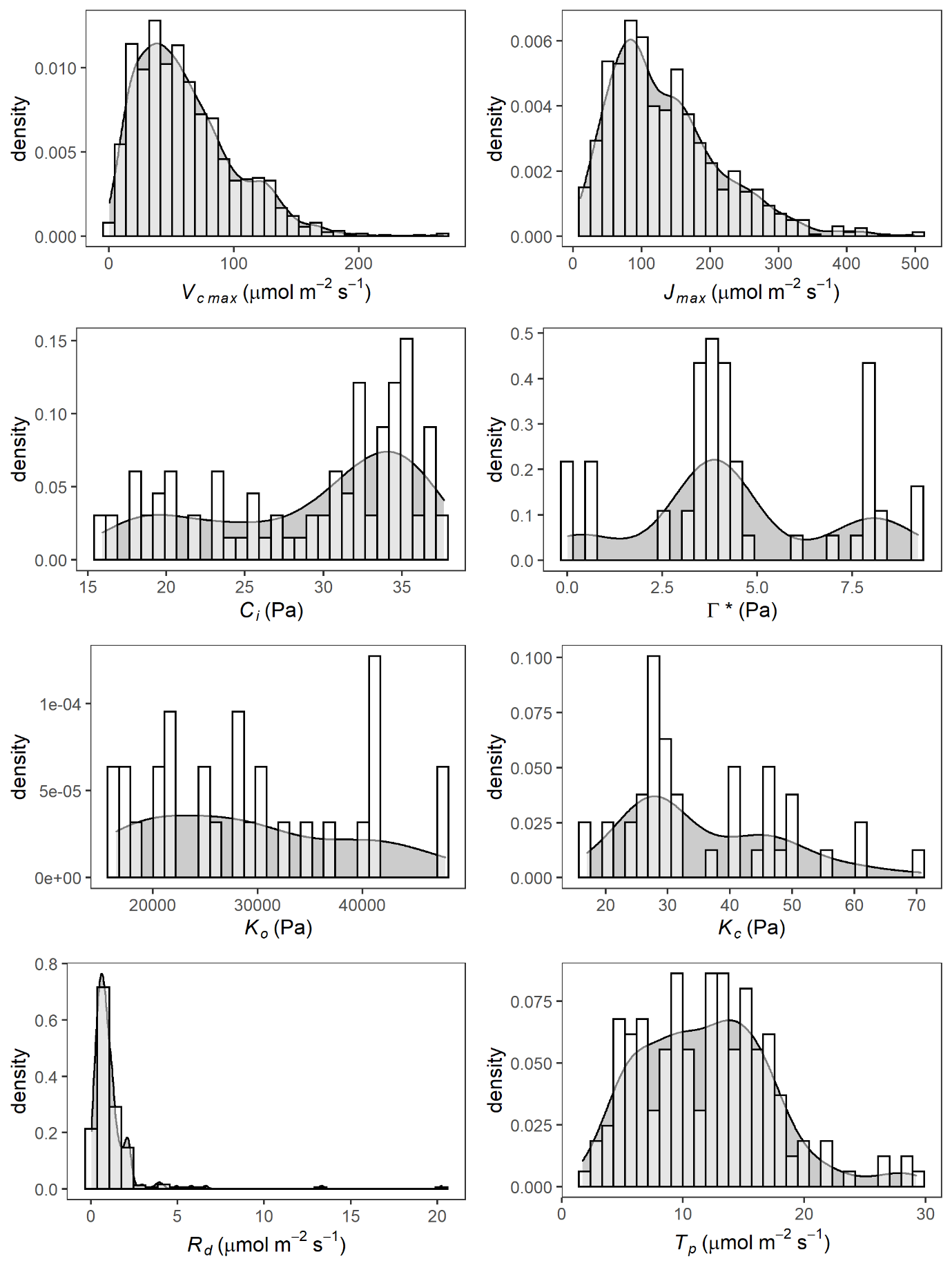

**Fig S1**: Global probability density distributions for each parameter of the Farquhar photosynthesis equations 1-3, 6 and 7 across all species for which data were available. Note that not every study estimated every parameter, and so the sample size varies across each parameter. Summary statistics of all parameters are shown in Table 2, parameter definitions and descriptions in Table 1.

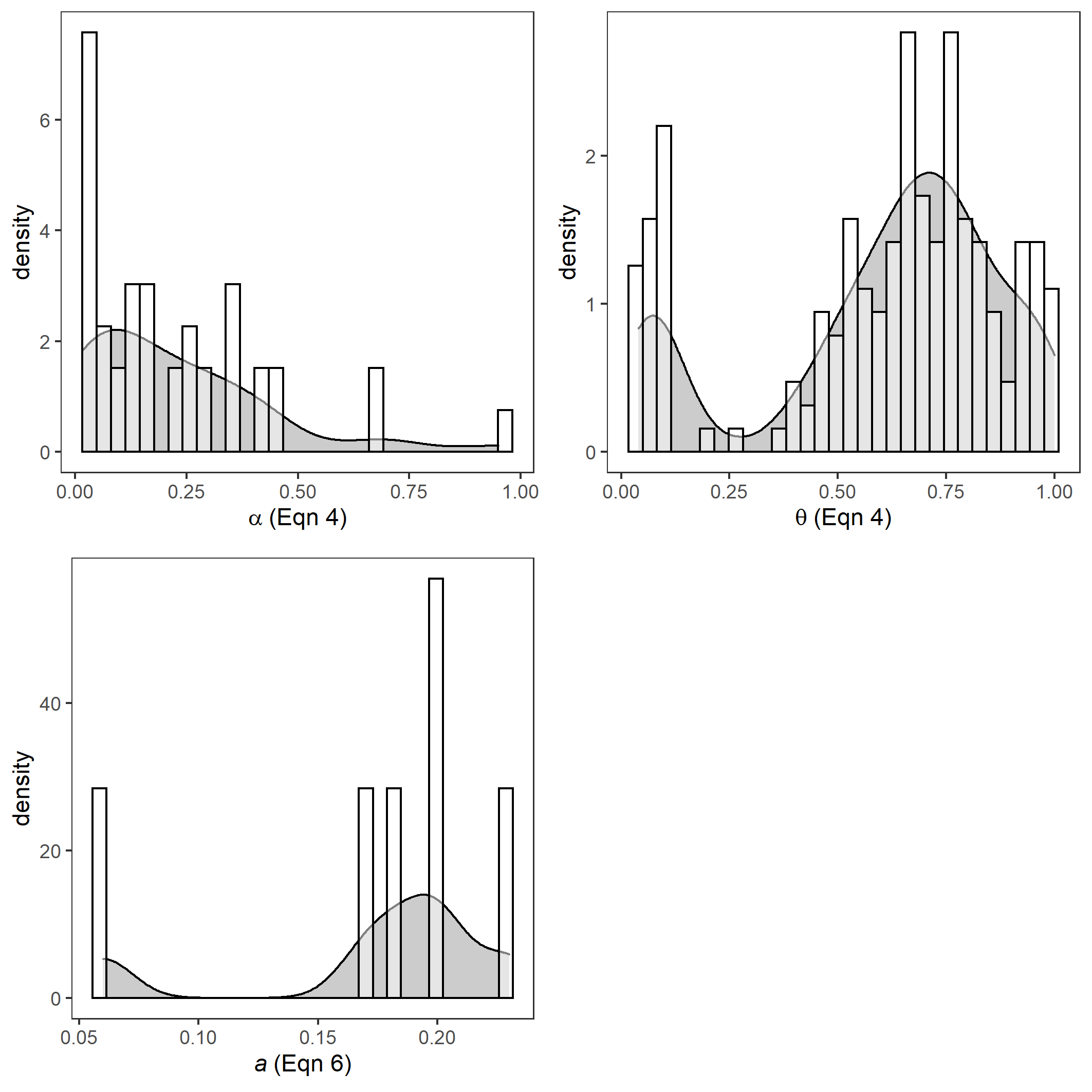

**Fig S2:** Global probability density distributions for each parameter of the Farquhar photosynthesis equations 4-5 across all species for which data were available. Note that not every study estimated every parameter, and so the sample size varies across each parameter. Summary statistics of all parameters are shown in Table 3, parameter definitions and descriptions in Table 1.

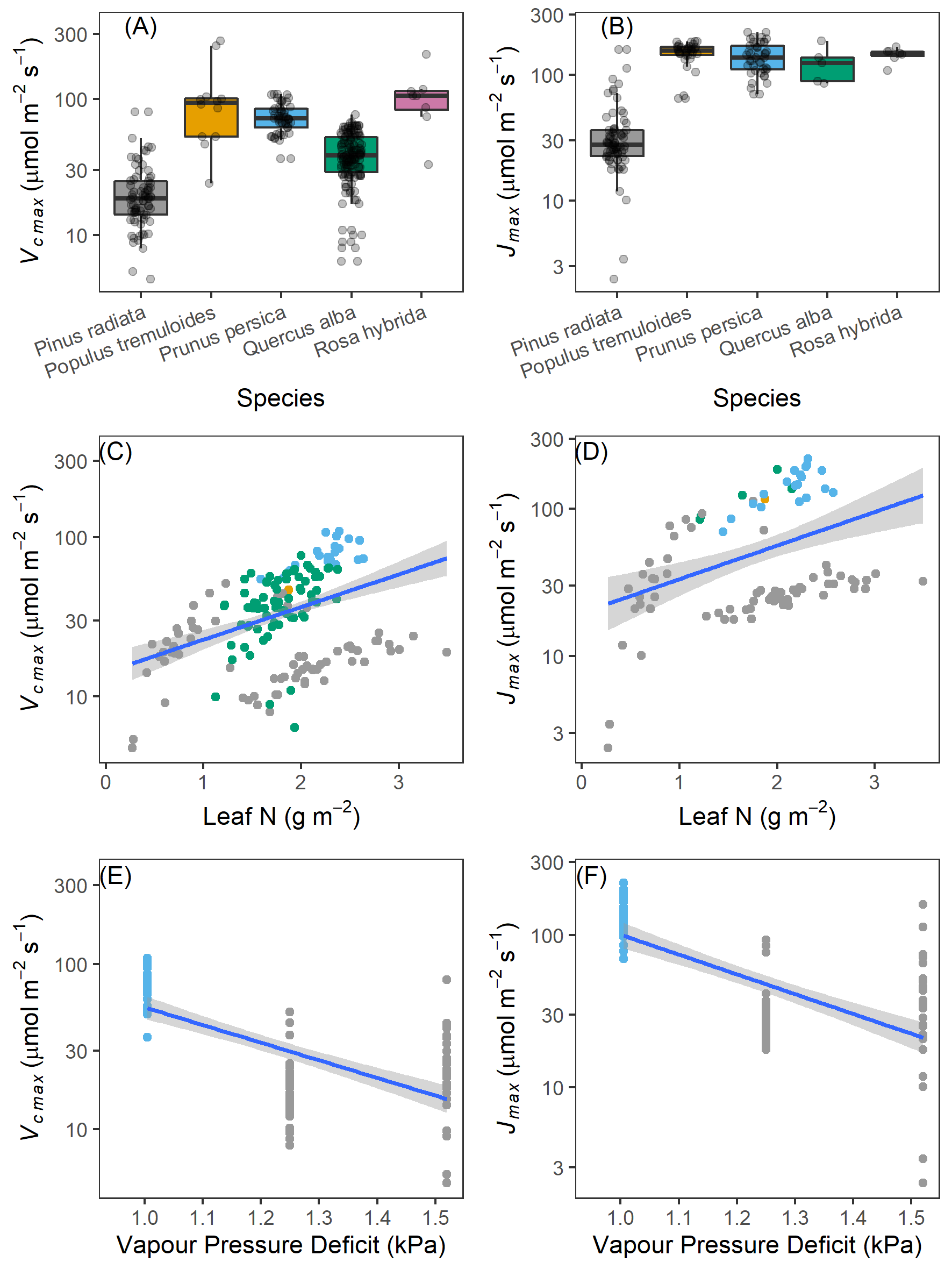

**FIGURE S3**: Comparison of variation within species for five species for which 10 or more different studies (not parameter estimates) existed for (A) $V_{c,max}$ or (B) $J_{max}$. Variation within a species can be over an order of magnitude. This variation however, can largely be explained by methodological differences such as (C, D) leaf temperature, or (E, F) leaf nitrogen content per area. Different species are coloured the same across all panels. The y-axis on panels C-F are on a log_10_-scale.

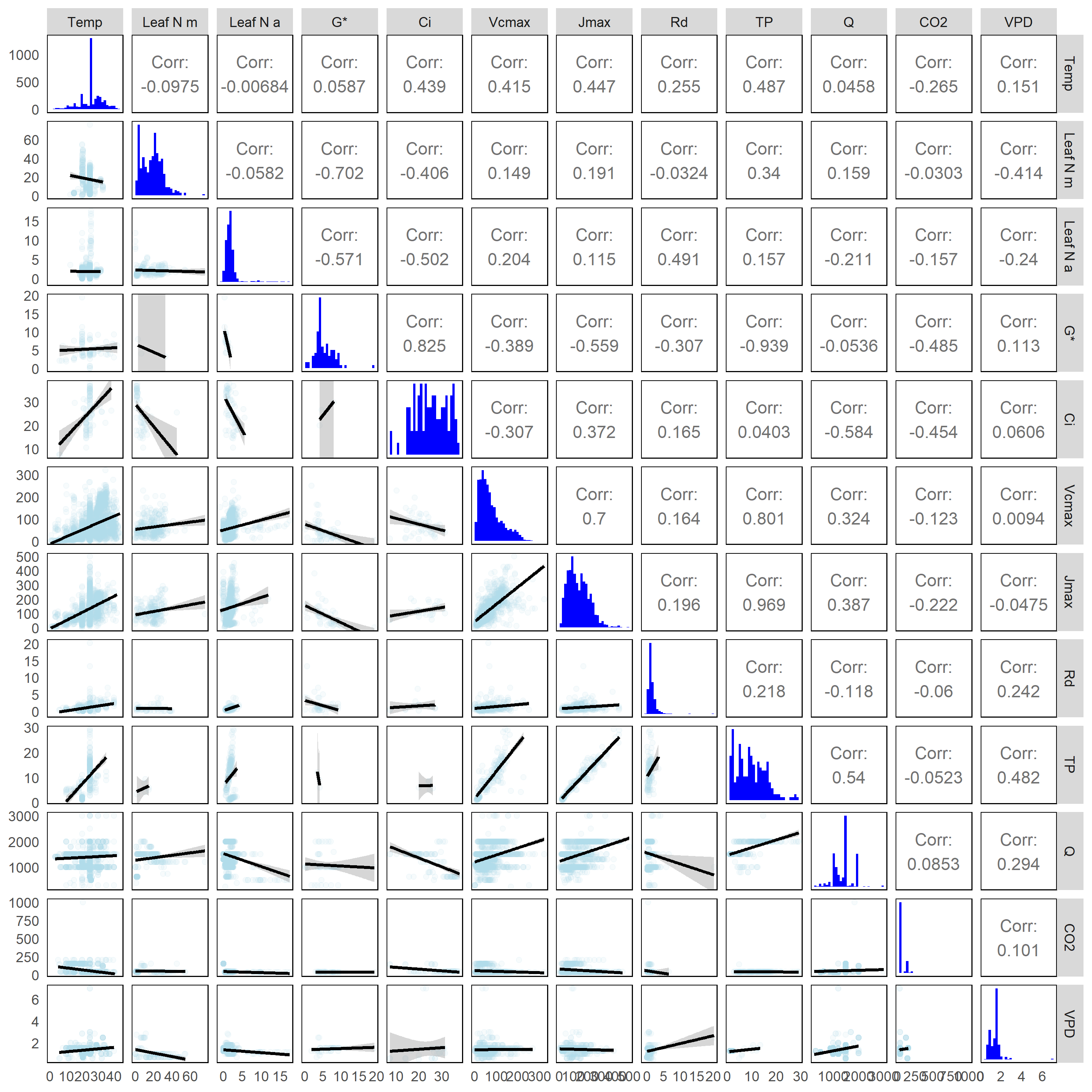

**FIGURE S4**: The variables and parameters used in the PCA analysis of the main text (Fig 4). The upper triangle matrix shows the Pearson correlation coefficient of each variable, while the lower triangle matrix shows the associated bivariate scatter plot (blue) including a regression line (black) and its confidence interval (gray). The diagonal shows the density histogram of each variable. Parameter labels are as listed in Table 1. Variables are: Temp=temperature; Q=irradiance used for $A-C$ curve; Leaf N m = leaf nitrogen content per mass; Leaf N a = leaf nitrogen content per area; VPD = vapour pressure deficit, and; CO2 = CO_2_ level used for $A-Q$ curve. Though we had more variables than this (E.g. LMA and SLA), there were not enough data to estimate correlations for variables not shown here.

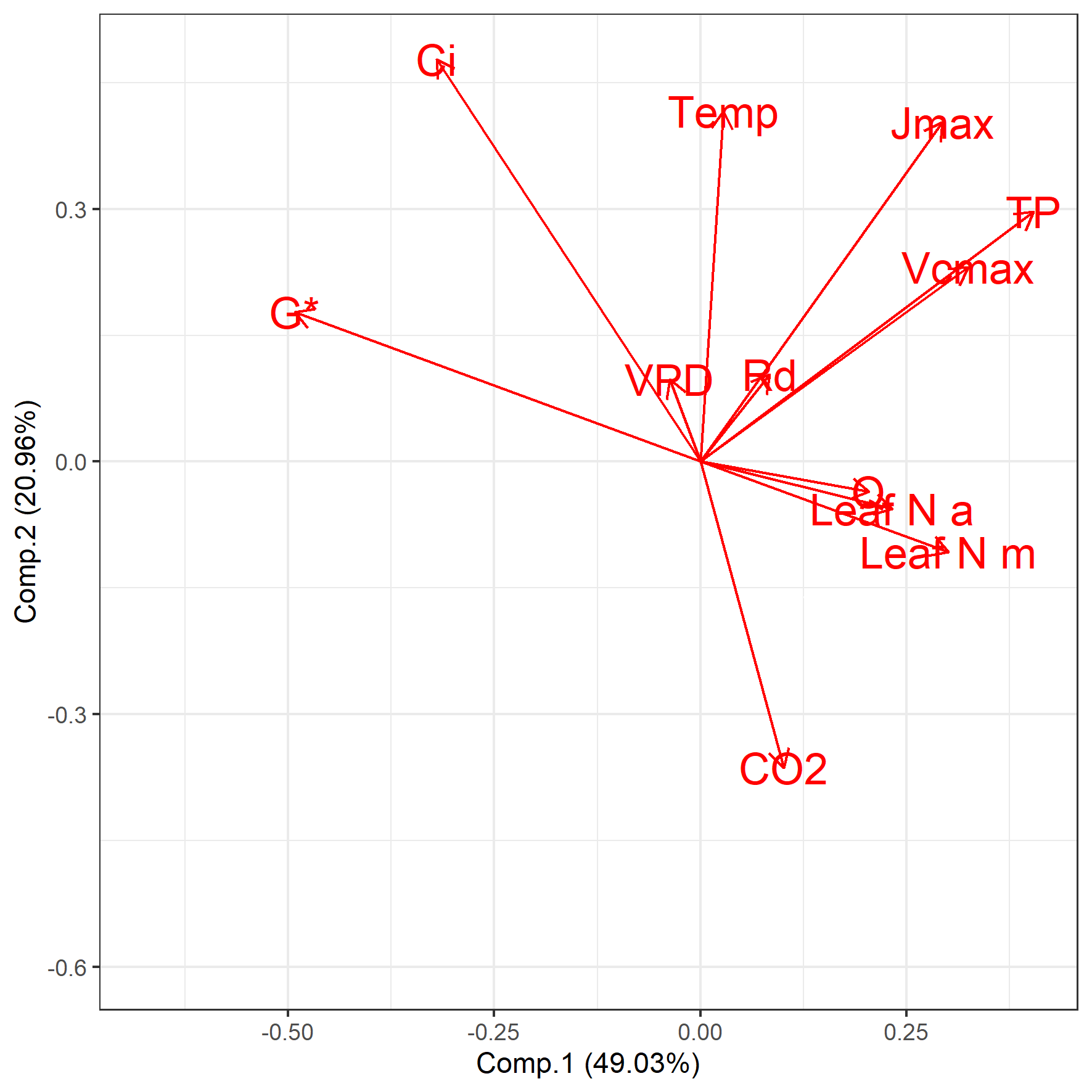

**FIGURE S5**: Results of the principle components analysis of the correlation matrix for which pairwise complete observations existed. Vectors pointing in the same direction are positively correlated, but the strength of the correlation decreases as the angle between vectors increases, becoming 0 at 90°. Thus, vectors at right angles are orthogonal, and uncorrelated. Vectors pointing in opposite directions are negatively correlated, and the strength of this correlation decreases as the angle between vectors decreases, becoming 0 at 90°. Pairwise bivariate plots of all variables are shown in Fig S3. Parameter vector labels are as listed in Table 1. Other variables vector labels are: Temp=temperature; Q=irradiance used for $A-C_{i}$ curve; Leaf N m = leaf nitrogen content per mass; Leaf N a = leaf nitrogen content per area; VPD = vapour pressure deficit, and; CO2 = CO_2_ level used for $A-Q$ curve.

**The phylogeny from Figure 1 in NEWICK code format.**

(((((((((((Polygonum_viviparum:54.0144,Polygonum_cuspidatum:54.0144,Polygonum_lapathifolium:54.0144,Rumex_alpestris:54.0144,Rumex_verticillatus:54.0144,Polygonum_pensylvanicum:54.0144)Polygonaceae:29.3923,Tamarix_ramosissima:83.4067):6.05427,(Simmondsia_chinensis:83.2984,((Beta_vulgaris:28.9453,(Spinacia_oleracea:22.0065,Chenopodium_album:22.0065):6.9388):0.75562,Haloxylon_ammodendron:29.7009):53.5975):6.16252)Caryophyllales:27.2363,(((((Vaccinium_darrowii:62.0517,(Arbutus_unedo:53.6508,(Calluna_vulgaris:43.5038,(Oxydendrum_arboreum:24.104,((Vaccinium_myrtillus:7.76228,(Vaccinium_corymbosum:6.8366,Vaccinium_bracteatum:6.8366):0.92568):3.62938,Vaccinium_uliginosum:11.3917):12.7124):19.3998):10.147):8.40083)Ericaceae:20.7165,(Eurya_japonica:71.8252,Schima_superba:71.8252):10.943):2.80459,Aegiceras_corniculatum:85.5727):20.1031,Nyssa_aquatica:105.676):1.93366,(((((Artemisia_giraldii:45.7437,Lactuca_saliva:45.7437,Dendrosenecio_keniodendron:45.7437,Helianthus_peliolaris:45.7437,Eupatorium_maculatum:45.7437,((Adenocaulon_bicolor:15.0349,Cynara_cardunculus:15.0349):19.9056,(Senecio_vulgaris:28.2725,(((Artemisia_tridentata:3.58929,(Artemisia_gmelinii:2.52209,Artemisia_campestris:2.52209):1.0672):1.04717,Achillea_millefolium:4.63646):19.9573,((Mikania_micrantha:2.05431,Mikania_cordata:2.05431):21.7388,((Bidens_frondosa:2.90989,Bidens_cernua:2.90989):16.8747,(Arnica_montana:13.4546,(Helianthus_annuus:11.3008,(Ambrosia_artemisiifolia:1.73541,Xanthium_strumarium:1.73541):9.5654):2.15375):6.33002):4.00852):0.800646):3.67877):6.66798):10.8032)Asteraceae:37.92,(Lobelia_lelekii:52.9672,(Cyanea_hirtella:35.8155,Campanula_scheuchzeri:35.8155):17.1518)Campanulaceae:30.6964)Asterales:11.7531,(Brassaia_actinophylla:76.5788,Sium_suave:76.5788):18.838):5.57251,Ilex_pedunculosa:100.989)Campanulidae:1.88221,(((((Capsicum_annum:47.0276,Lycopersicon_esculenlum:47.0276,Lycopersicon_esculentum:47.0276,Nicotiana_tobaccum:47.0276,((Nicotiana_sylvestris:4.00367,Nicotiana_tabacum:4.00367):10.15,(Capsicum_annuum:6.8691,(Solanum_lycopersicum:1.13229,Solanum_tuberosum:1.13229):5.73681):7.2846):32.8739)Solanaceae:9.21285,Pharbitis_nil:56.2405):16.5865,(((((Lycopus_americanus:41.3946,(Tabebuia_rosea:41.0622,Scrophularia_desertorum:41.0622):0.332342):0.621584,Avicennia_marina:42.0162):0.129732,Rhinanthus_alectorolophus:42.1459):0.615055,(Plantago_atrata:4.11564,(Plantago_maritima:2.93056,(Plantago_major:0.991475,Plantago_media:0.991475):1.93909):1.18507):38.6453):13.4796,(Ligustrum_japonica:39.3526,Olea_europea:39.3526,(Phillyrea_angustifolia:22.751,(Fraxinus_pennsylvanica:16.8991,Fraxinus_excelsior:16.8991):5.85189):16.6016)Oleaceae:16.8879):16.5865):0.542387,(Nerium_oleander:58.8049,Nauclea_diderrichii:58.8049)Gentianales:14.5645):23.5808,Aucuba_japonica:96.9503):5.92122):4.73798)Asteridae:9.08781)Pentapetalae:2.46823,((((((((((((Gleichenia_squamulosa:0.397811,Pteris_vitatta:0.397811,Quintinia_serrata:0.397811,Hypolepis_poeppigii:0.397811,Athyrium_filixfemina:0.397811,Larix_decidua_x_leptolepis:0.397811,(Dryopteris_filix-mas:0.265207,Dryopteris_tyrrhena:0.265207,Polistichium_aculeatum:0.265207)dryopteridaceae:0.132604,Thelypteris_dentata:0.397811,(Blechnum_gibbum:0.265207,Blechnum_magallanicum:0.265207,Blechnum_mochaenum:0.265207,Blechnum_pennamarina:0.265207)blechnaceae:0.132604,Chenopodium_polyspermum_:0.397811,Lophosoria_quadripinnata:0.397811,Nephrolepsis_exaltata:0.397811)Abutilon_theophrasti:18.2921,(Gossypium_hirsutum:2.35364,Gossypium_barbadense:2.35364):16.3362):1.08918,Hibiscus_cannabinus:19.7791):49.3403,Argyrodendron_trifoliolalum:69.1194,Malvastrum_rotundifolium:69.1194,Gossypium_hirsulum:69.1194,Gossypium_hirsulwn:69.1194)Malvaceae:10.6293,(Cistus_salvifolius:49.1673,(Dipterocarpus_glabosus:40.9998,Dryobalanops_aromatica:40.9998,(Shorea_beccariana:0.043711,(Shorea_macroptera:0.030878,Shorea_acuta:0.030878):0.012833):40.9561)Dipterocarpaceae:8.16752):30.5814):20.7649,((((Toona_ausiralis:52.7612,Entandrophragma_angolense:52.7612)Meliaceae:0.821294,((Citrus_limon:1.54686,Citrus_sinensis:1.54686):27.0242,Flindersia_brayleyana:28.5711):25.0114):15.0801,(Dodoea_triquetra:66.2502,Dodonaea_angustissima:66.2502,(Acer_rubrum:10.0337,(Acer_pseudoplatanus:7.61713,Acer_saccharum:7.61713):2.41659):56.2165)Sapindaceae:2.41233):2.27819,(Anacardium_occidenlale:59.0955,Pistacia_lentiscus:59.0955)Anacardiaceae:11.8453):29.5729):0.908691,(Arabidopsis_thaliana:14.0671,((Brassica_rapa:0.983766,(Brassica_napus:0.942695,Brassica_oleracea:0.942696):0.041071):0.131126,(Raphanus_sativus:0.880426,Diplotaxis_erucoides:0.880427):0.234465):12.9522):87.3552):9.59952,((((Corymbia_latifolia:71.3021,Eucalyptus_miniata:71.3021,Eucalyptus_pruinosa:71.3021,Eucalyptus_tectifica:71.3021,(Metrosideros_umbellata:44.6971,((Leptospermum_scoparium:12.4057,Kunzea_ericoides:12.4057):23.9573,((Corymbia_aparrerinja:12.7745,Corymbia_terminalis:12.7745):20.5061,(Eucalyptus_tetrodonta:29.2182,(Eucalyptus_pauciflora:19.3526,(Eucalyptus_coolabah:7.98054,(Eucalyptus_grandis:4.98689,Eucalyptus_globulus:4.98689):2.99365):11.3721):9.86557):4.0625):3.08232):8.33415):26.6049)Myrtaceae:15.1908,Lythrum_salicaria:86.4929):0.728903,(Combretum_nigricans:65.4722,(Guiera_senegalensis:31.2241,Combretum_micranthum:31.2241):34.2482)Combretaceae:21.7496):0.978722,Fuchsia_excorticata:88.2005)Myrtales:22.8213):2.22862,Geranium_sylvaticum:113.25)Malvidae:3.62701,Vitis_vinifera:116.877):0.657279,(((((((Populus_delloides:78.4744,Populus_euramericana:78.4744,Populus_fremontii:78.4744,Populus_grandidenlata:78.4744,Populus_x_euramericana:78.4744,((Populus_euphratica:25.6223,(Populus_deltoides:17.1672,(Populus_balsamifera:11.972,(Populus_nigra:8.85335,(Populus_grandidentata:6.4343,Populus_tremuloides:6.4343):2.41905):3.11868):5.19513):8.45518):8.01784,Salix_dasyclados:33.6402):44.8342)Salicaceae:22.1069,(Manihot_esculentum:99.388,Claoxylon_sandwicense:99.388,Ricinus_communis:99.388)Euphorbiaceae:1.19323):0.532812,Triadenum_fraseri:101.114):4.73399,Goupia_glabra:105.848)Malpighiales:5.53298,(Weinmannia_racemosa:82.5775,Aristotelia_serrata:82.5775):28.8035):1.87759,((Glycine_wightii:69.1425,Vigna_unguiculala:69.1425,Caragana_korshinskii_:69.1425,Phaseolus_atropurpureus:69.1425,Acacia_mangium:69.1425,Amphimas_pterocarpoides:69.1425,(Acacia_ligulata:57.5823,((Lespedeza_davurica:21.2758,((Vigna_luteola:3.19408,(Phaseolus_acutifolius:0.412871,Phaseolus_vulgaris:0.412871):2.78121):3.34919,(Calopogonium_mucunoides:3.17134,Glycine_max:3.17134):3.37193):14.7325):20.4598,(Robinia_pseudoacacia:35.2593,(Cicer_arietinum:18.1933,(Vicia_faba:14.1538,(Trifolium_alpinum:9.63646,(Trifolium_pratense:4.53159,(Trifolium_montanum:2.20199,Trifolium_repens:2.20199):2.3296):5.10487):4.51734):4.03951):17.066):6.47628):15.8467):11.5602)Fabaceae:37.8529,((((Quercus_petrea:40.41,(Fagus_grandifolia:33.9289,(Fagus_sylvatica:19.0026,Fagus_crenata:19.0026):14.9263):6.48114,((Castanea_sativa:2.12713,Castanopsis_cuspidata:2.12713):2.41037,((Quercus_glauca:1.56808,(Quercus_ilex:1.524,Quercus_suber:1.524):0.04409):1.20986,((Quercus_myrtifolia:1.21586,(Quercus_velutina:0.127739,Quercus_rubra:0.127739):1.08812):0.853149,(Quercus_petraea:1.33892,(Quercus_robur:0.291445,((Quercus_pyrenaica:0.038691,Quercus_serrata:0.038691):0.109713,(Quercus_geminata:0.078292,(Quercus_stellata:0.059289,Quercus_alba:0.059289):0.019003):0.070112):0.14304):1.04748):0.730089):0.708931):1.75955):35.8725)Fagaceae:21.8954,(((Carya_ovata:13.5678,Carya_glabra:13.5678):10.7427,(Juglans_regia:10.551,Juglans_nigra:10.551):13.7595):25.063,(Betula_alleghaniensis:14.3659,(Betula_papyrifera:3.91551,Betula_pendula:3.91551):10.4504):35.0075):12.9319):40.7694,Cucumis_sativus:103.075):2.86119,((Rosa_hybrida:82.294,Malus_domestica:82.294,Prunus_cerasus:82.294,Prunus_x_yedoensis:82.294,Heteromeles_arbulifolia:82.294,(Alchemilla_vulgaris:46.1879,(Duchesnea_indica:26.2426,Potentilla_aurea:26.2426):19.9453):36.1061,(Purshia_tridentata:81.4337,((Prunus_serotina:23.2159,Prunus_persica:23.2159):3.01031,Malus_pumila:26.2262):55.2075):0.860363)Rosaceae:6.98304,(Ceanothus_megacarpus:79.5238,((Celtis_adolfi-friderici:65.4822,Ficus_obtusifolia:65.4822):8.75071,(Musanga_cecropioides:63.2396,Hippophae_rhamnoides:63.2396):10.9932):5.29088):9.75332)Rosales:16.6589):1.05941):6.26322)Fabidae:0.420986,(Larrea_divaricaia:72.8237,Larrea_tridentata:72.8237)Zygophyllaceae:40.8559):3.85514)Rosidae:0.138911,(Daphniphyllum_humile:107.877,(Liquidambar_styraciflua:105.745,Cercidiphyllum_japonicum:105.745):2.13196):9.79685)Superrosidae:1.49186)Gunneridae:17.735,((Pulsatilla_sulphurea:105.446,Anemone_raddeana:105.446,(Trollius_europaeus:69.1438,Ranunculus_acris:69.1438):36.3018)Ranunculaceae:30.6465,(((Hakea_tephrosperma:92.3053,(Leucadendron_xanthoconus:0.957953,Leucadendron_laureolum:0.957954):91.3474)Proteaceae:2.50544,Platanus_orientalis:94.8108):39.5967,Pachysandra_terminalis:134.407):1.68465):0.808344)Eudicotyledoneae:43.7278,((Piper_hispidum:9.57395,Piper_auritum:9.57395):137.491,((Cinnamomum_camphora:34.7772,Sassafras_albidum:34.7772):85.9232,Liriodendron_tulipifera:120.7):26.3648)Magnoliidae:33.5631):7.65496,(((((Glyceria_canadensis:47.7638,Triticum_aeslivum:47.7638,Digitaria_hispidula:47.7638,Poa_compressea:47.7638,Sesleria_varia:47.7638,Agrostis_tenuis:47.7638,Hordeum_glaucum:47.7638,Hordeum_vulgar:47.7638,Digitaria_ischaemum:47.7638,Oryza_saliva:47.7638,Pharus_latifolius:47.7638,((Nardus_stricta:8.4423,((Hordeum_vulgare:3.10189,Triticum_aestivum:3.10189):1.02103,((Dactylis_glomerata:2.14617,((Festuca_arundinacea:1.11513,(Lolium_multiflorum:0.294414,Lolium_perenne:0.294414):0.820712):0.82964,Festuca_rubra:1.94477):0.201408):0.690797,(Avenella_flexuosa:2.46532,((Phalaris_arundinacea:1.44754,(((Trisetum_flavescens:0.551699,Koeleria_pyramidata:0.5517):0.550165,Avena_sativa:1.10186):0.119316,(Briza_media:0.807936,Agrostis_capillaris:0.807936):0.413244):0.22636):0.817689,Poa_trivialis:2.26523):0.20009):0.371652):1.28594):4.31939):10.6863,((Leersia_oryzoides:5.35536,Oryza_sativa:5.35535):12.6149,((Bothriochloa_ischaemum:5.70535,Cenchrus_ciliaris:5.70535):3.97523,(Spartina_alterniflora:2.94396,Eragrostis_pectinacea:2.94397):6.73662):8.28967):1.15836):28.6352)Poaceae:36.4567,(Carex_projecta:8.08412,((Carex_folliculata:4.37601,Carex_crinita:4.37601):2.2784,(Carex_retrorsa:0.823806,Carex_tuckermanii:0.823807):5.8306):1.42971):76.1364):38.951,(Haumania_danckelmaniana:59.2463,Megaphrynium_macrostachyum:59.2463)Marantaceae:63.9252)Commelinidae:11.6939,(Dasylirion_leiophyllum:97.875,Iris_versicolor:97.875):36.9904):23.4767,(Alocasia_macrorrhiza:123.182,Colocasia_esculent:123.182)Araceae:35.1604)Nartheciidae:29.9412):163.951,((Tsuga_canadensis:237.575,((Larix_gmelinii:174.927,Pseudotsuga_menziesii:174.927):32.6935,((Picea_sitchensis:171.666,((Picea_glauca:3.73968,Picea_engelmannii:3.73968):79.9448,((Picea_rubens:46.059,Picea_mariana:46.059):13.7487,Picea_abies:59.8076):23.8769):87.9814):12.8177,((Pinus_strobus:58.5776,Pinus_flexilis:58.5776):93.5894,(Pinus_banksiana:143.836,((Pinus_pinaster:63.9799,(Pinus_sylvestris:2.45928,Pinus_densiflora:2.45928):61.5206):18.8999,(Pinus_contorta:72.9023,(Pinus_ponderosa:68.5124,(Pinus_taeda:58.7314,Pinus_radiata:58.7314):9.78097):4.38987):9.97752):60.9564):8.33082):32.3167):23.1367):29.9543)Pinales:29.1416,(Chamaecyparis_obtusa:230.88,(Prumnopitys_ferruginea:169.862,Dacrydium_cupressinum:169.862)Podocarpaceae:61.0179)Cupressophyta:35.8365):85.5185)Spermatophyta:38.4679,((Botrychium_lunaria:138.034,Ophioglossum_vulgatum:138.034):191.959,(((Equisetum_telmateia:146.11,(Equisetum_sylvaticum:22.3197,Equisetum_pratense:22.3197):123.79):32.0246,Equisetum_arvense:178.135):126.099,(Osmunda_regalis:239.283,(Lygodium_japonicum:157.507,(Adiantum_pedatum:118.841,(Pteridium_aquilinum:104.219,((Asplenium_scolopendrium:14.1064,Asplenium_trichomanes:14.1064):76.5276,(((Gymnocarpium_dryopteris:53.6437,Cystopteris_sudetica:53.6437):15.1439,(Thelypteris_palustris:46.6804,Matteuccia_struthiopteris:46.6804):22.1072):8.99504,(Dryopteris_erythrosora:24.5278,Dryopteris_carthusiana:24.5278):53.2548):12.8514):13.5853):14.6217):38.6664):81.775):64.9515):25.7593):60.7092):10.0851,(Lycopodium_clavatum:55.5091,Lycopodium_annotinum:55.5091):345.279);
